# Heat-evolved algal symbionts enhance bleaching tolerance of adult corals without trade-off against growth

**DOI:** 10.1101/2023.06.17.545444

**Authors:** Wing Yan Chan, David Rudd, Luka Meyers, Sanjida H. Topa, Madeleine J. H. van Oppen

## Abstract

Ocean warming has caused coral mass bleaching and mortality worldwide and the persistence of symbiotic reef-building corals requires rapid acclimation or adaptation. Experimental evolution of the coral’s microalgal symbionts followed by their introduction into coral is one potential method to enhance coral thermotolerance. Heat-evolved microalgal symbionts of the generalist species, *Cladocopium proliferum* (strain SS8), were exposed to elevated temperature (31°C) for ∼10 years, and were introduced into chemically bleached adult fragments of the scleractinian coral, *Galaxea fascicularis*. The new symbionts persisted for the five months of the experiment and enhanced adult coral thermotolerance compared with corals that were inoculated with the wild-type *C. proliferum* strain. Thermotolerance of SS8-corals was similar to that of coral fragments from the same colony hosting the homologous symbiont, *Durusdinium* sp., which is naturally heat-tolerant. However, SS8-coral fragments exhibited faster growth and recovered cell density and photochemical efficiency more quickly following chemical bleaching and inoculation under ambient temperature relative to *Durusdinium*-corals. Mass spectrometry imaging suggests that algal pigments involved in photobiology and oxidative stress were the greatest contributors to the thermotolerance differences between coral hosting heat-evolved versus wild-type *C. proliferum*. These pigments may have increased photoprotection in the heat-evolved symbionts. Our findings show that adult coral thermotolerance can be enhanced via the uptake of exogenously supplied, heat-evolved symbionts, without a trade-off against growth under ambient temperature. Heat-evolved *C. proliferum* remains in the corals in moderate abundance two years after its first inoculation, suggesting long-term stability of this novel symbiosis.

## 1. INTRODUCTION

Tropical coral reefs are among the most biodiverse ecosystems on Earth. The survival of reef-building corals depends on nutrient exchange between the coral host and its microalgal photosymbionts (family: Symbiodiniaceae) (Davy et al. 2012). These corals cannot survive without the carbon and nitrogen translocated from Symbiodiniaceae (Muller-Parker et al. 2015), and symbiodiniacean physiological properties largely determine their coral host’s thermotolerance (Berkelmans and van Oppen 2006; Quigley et al. 2018; Palacio-Castro et al. 2023). However, climate change increases sea surface temperatures and the frequency, duration and intensity of marine heat waves (Bindoff et al. 2019), causing the dissociation of this obligate partnership (i.e., coral bleaching) that results in coral starvation and often death (Hoegh-Guldberg 1999). Globally, marine heat waves are linked to coral population decline in the last two to three decades (Hughes et al. 2018).

Given the significance of the coral-Symbiodiniaceae symbiosis, introducing heat-tolerant Symbiodiniaceae into corals is one potential way to enhance coral thermotolerance and maximize the likelihood of coral survival under ocean warming. Many members of the symbiodiniacean genus *Durusdinium* are naturally heat-tolerant and are often found in higher abundance in corals from extreme or fluctuating environments, or after a mass bleaching event (Baker et al. 2004; Oliver and Palumbi 2011; Stat and Gates 2011; Boulotte et al. 2016; Camp et al. 2019; Quigley et al. 2022; Palacio-Castro et al. 2023). Higher thermotolerance is generally seen in corals hosting *Durusdinium* sp. when compared to those hosting members of the genus *Cladocopium* (Berkelmans and van Oppen 2006; Silverstein et al. 2015; Palacio-Castro et al. 2023) – the most widespread symbiont genus in Indo-Pacific scleractinian corals (LaJeunesse 2005). However, the physiological trade-offs often associated with *Durusdinium* sp. under ambient temperatures make it a less ideal candidate for reef restoration (Ortiz et al. 2013). Trade-offs include less photosynthate translocation to the host (Cantin et al. 2009; Baker et al. 2013; Allen-Waller and Barott 2023), as well as slower coral growth (Little et al. 2004; Cunning et al. 2015), lower amounts of stored lipids and smaller eggs (Jones and Berkelmans 2011) when compared to conspecific coral hosting *Cladocopium*. For instance, coral juveniles harbouring *Cladocopium* C1 had double the amount of ^14^C photosynthate (Cantin et al. 2009), 22% more ^15^N acquisition (Baker et al. 2013) and higher relative electron transport rate of photosystem II (Cantin et al. 2009) than those dominated by *Durusdinium* sp..

Experimental evolution has successfully enhanced the thermotolerance of multiple marine microalgal species (Chan et al. 2021), including *Cladocopium proliferum* (Chakravarti et al. 2017; Butler et al. 2023). Some heat-evolved *C. proliferum* strains also enhanced coral bleaching tolerance when in symbiosis with larvae (Buerger et al. 2020) or recruits (Quigley and van Oppen 2022). While these findings are promising, it is unclear whether they can be extrapolated to the adult life phase given coral larvae and adults differ significantly physiologically. Further, in contrast to coral larvae which are often aposymbiotic, adult corals cannot be rendered fully aposymbiotic with thermal (Coffroth et al. 2010) or chemical bleaching (Wang et al. 2012; Scharfenstein et al. 2022; Puntin et al. 2023). It is not known whether heat-evolved Symbiodiniaceae can establish a population in adult corals in the presence of remnant native Symbiodiniaceae as potential competitors.

This study tests whether heat-evolved *C. proliferum* can form a symbiosis with adult corals and if so, whether the novel symbiosis can enhance adult coral bleaching tolerance. The coral *G. fascicularis* was chosen, given it is known to host both *Cladocopium* (C1, C21a) and *Durusdinium* (D1-4) naturally (Wepfer et al. 2020; Puntin et al. 2023). The GBR *G. fascicularis* colonies of this study host *Durusdinium trenchii*, *Durusdinium glynnii*, and/or *Cladocopium* C40 in the wild (see Results). We removed > 99% of the coral’s native Symbiodiniaceae with chemical bleaching and inoculated them three times with 1) a strain of *C. proliferum* (SS8) exposed to 31°C in the laboratory for ∼10 years (SS8-corals), 2) a wild-type *C. proliferum* strain derived from the same mother culture exposed to 27°C for ∼10 years (WT10-corals), or 3) the homologous Symbiodiniaceae freshly isolated from *G. fascicularis* (a mix of *D. trenchii*, *D. glynnii* and C40) (HOM-corals). Control corals with no menthol treatment nor inoculation were also included; the Symbiodiniaceae communities of these corals were dominated by either *D. trenchii* and *D. glynnii* (Native-corals) or *Cladocopium* C40 (NativeC-corals). Eighteen weeks after the first inoculation, these holobionts were subjected to a simulated short-term heat stress event (32°C, 8 days) and their Symbiodiniaceae community composition, Symbiodiniaceae cell density, Symbiodiniaceae photochemical efficiency, coral size, coral survival, and spatial metabolite profiles were assessed. Note that NativeC-corals were only used for metabolite profiling due to low number of replicates. Metabolite profiles of the holobionts were revealed via high-resolution mass spectrometry imaging (MSI), which can provide spatial and relative intensity information on a wide range of metabolites with minimum biomass requirement (Chan et al. 2023).

## 2. MATERIALS AND METHODS

### 2.1 Coral inoculation experiment

#### 2.1.1 Coral colonies

Experimental timeline is shown in Fig. 1 and Table S1. Five colonies of *Galaxea fascicularis* (genotype G1-G5) were obtained from Falcon Reef, the GBR (18°45.989’S, 146°32.130’E) on 29^th^ Oct 2020 and transported by air in seawater-filled bags in polystyrene boxes to the School of BioSciences, the University of Melbourne. All colonies had a similar polyp size (∼1 cm diameter per polyp) and were maintained at 27°C in a 120 L recirculating aquarium system with reconstituted seawater made from Red Sea Salt and reverse osmosis water. Coral colonies were cut into ∼4-polyp fragments (*n* = 40-50 per genotype) with a diamond band saw (Gryphon C40 CR Custom XL) and secured onto a 22 mm diameter ceramic plug (Seachem Reef Glue^TM^) (Fig. 1; Table S2). The fragments were returned to the aquaria for a 10-day recovery period to allow the cut surfaces to be healed prior to menthol bleaching. The remaining part of the colonies were kept in the recirculating system to obtain homologous Symbiodiniaceae for inoculation. The corals were maintained under a diurnal light profile with ∼130-155 µE m^-2^ s^-1^ at full sun (14:10h light: dark) and were fed with a mix of microalgae (Shellfish Diet 1800®) at 5000 cell mL^-1^ three times a week. A total of 30% of the seawater was changed weekly.

**Fig. 1.**
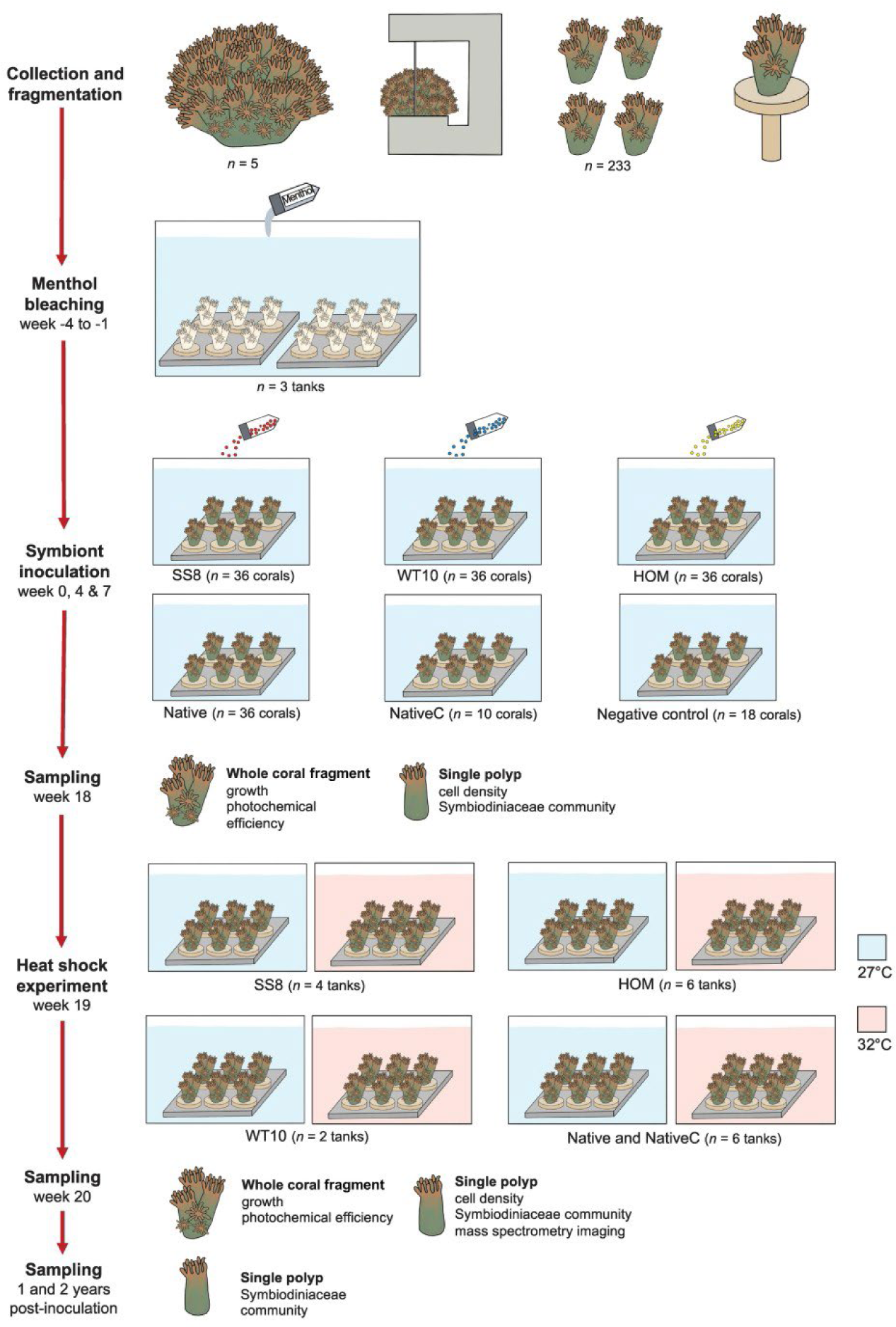
Simplified experimental design and timeline. The abbreviation of the inocula and coral groups refer to: heat-evolved *Cladocopium proliferum* (SS8-corals), wild-type *C. proliferum* (WT10-corals), freshly isolated, homologous Symbiodiniaceae (a mix of ∼80% *Durusdinium* sp. and ∼13% *Cladocopium* C40; see Results) from *Galaxea fascicularis* (HOM-corals), as well as control corals with no menthol treatment and no inoculation that were natively dominated by *Durusdinium trenchii* and *D. glynnii* (Native-corals) or *Cladocopium* C40 (NativeC-corals). Note that NativeC-corals were only used for mass spectrometry imaging due to their low number of replicates. See Table S1 for full details on the experimental timeline.

#### 2.1.2 Menthol bleaching and Symbiodiniaceae cell density

Menthol bleaching was used to remove the coral’s native Symbiodiniaceae population following our pilot study adapted from (Wang et al. 2012; Matthews et al. 2016; Puntin et al. 2023) (Supplementary Methods S1). Coral fragments were distributed randomly and evenly across three 9 L tanks and dosed with 0.38 mM of menthol for 8 h (08:30-16:30) under light. The recirculating pumps stayed on to ensure even mixing. After 8 h, the corals were removed from the tanks, rinsed in seawater and placed in the 120 L recirculating system to rest overnight (16:30-08:30). The procedure was repeated for four consecutive days and the corals were not fed. On the 5^th^ day, the corals were fed and left to rest in the 120 L system for three days. This weekly schedule was repeated for four weeks, until the corals were visually bleached as assessed under a high-power stereomicroscope (Leica M205 FA). A total of 46 coral fragments were not exposed to menthol treatment and were kept as control; of these, 36 fragments harboured Symbiodiniaceae communities dominated by *D. trenchii* and *D. glynnii* (Native-corals) and 10 had communities dominated by *Cladocopium* C40 (NativeC-coral) (Fig. 1).

To quantify the amount of remaining Symbiodiniaceae cells in the corals, five single polyps per genotype, with or without menthol treatment were sampled at the end of the four-week treatment (5 replicates x 4 genotypes x 2 treatments, *n* = 40). Sample tissues were removed by water-piking with 50 mL of filtered seawater (FSW), after which tissues were centrifuged at 3000 g for 5 min. The pellets were resuspended in 1.5 mL of FSW, centrifuged at 3000 g for 5 min and resuspend in 100 µL of FSW. Cell counts were conducted with a hemocytometer using 10 µL and two technical replicates per sample. A 20 times dilution with FSW was applied to control corals prior to cell counting.

#### 2.1.3 Symbiodiniaceae culture and inoculation

The chemically bleached corals were inoculated with three Symbiodiniaceae strains (hereafter referred to as Symbiodiniaceae treatments): heat-evolved *Cladocopium proliferum* (SS8), wild-type *C. proliferum* (WT10), and freshly isolated, homologous Symbiodiniaceae (a mix of ∼80% *Durusdinium trenchii*, *D. glynnii* and ∼13% *Cladocopium* C40; see Results) from *G. fascicularis* (HOM) (Table 1). The corals inoculated with the specific strains are referred to as “SS8-corals”, “WT10-corals”, or “HOM-corals”. SS8 was developed via experimental evolution (Chakravarti et al. 2017); it has better photochemical efficiency, growth rate and lower levels of extracellular reactive oxygen species than WT10 under elevated temperature of 31°C *in vitro* (Chakravarti et al. 2017). SS8 also conferred enhanced bleaching tolerant to *Acropora tenuis* coral larvae relative to WT10 (Buerger et al. 2020). Cultures of SS8 and WT10 were grown under their respective temperatures (Table 2) at 30-60 µE m^-2^ s^-1^ (12:12h, light: dark) in 1% IMK culture medium. The HOM inoculum was freshly isolated from a ∼5 cm diameter fragment of each *G. fascicularis* genotype. The Symbiodiniaceae were isolated via water-piking with ∼750 mL of FSW, and the blastate was centrifuged at 3000 g for 5 min and resuspended in FSW three times to remove residual coral tissue and mucus before coral inoculation.

**Table 1.** Details of the Symbiodiniaceae inocula of this study.

| **Symbiodiniaceae treatment** | **Species** | **Details** | **Culture ID** | **°C<sup>#</sup>** | **Original host** | **Host origin** |
| --- | --- | --- | --- | --- | --- | --- |
| WT10 | *Cladocopium proliferum* | Wild-type, cultured | SCF 055-01.10 | 27 | *Acropora tenuis* | Magnetic Island, Australia |
| SS8 | *Cladocopium proliferum* | Heat-evolved, conferring*, cultured | SCF 055-01.08 | 31 | *Acropora tenuis* | Magnetic Island, Australia |
| HOM | *Durusdinium trenchii*,<br>*D. glynnii* and<br>*Cladocopium* C40 | Wild-type, freshly isolated | NA | 27 | *Galaxea fascicularis* | Falcon Reef, Australia |

\* the heat-evolved strain SS8 was derived from the same mother culture as WT10. It was experimentally evolved under an elevated temperature of 31°C (Chakravarti et al. 2017) and it conferred enhanced bleaching tolerance to *Acropora tenuis* coral larvae (Buerger et al. 2020).
<sup>#</sup> indicates the temperature at which the cultures (WT10 and SS8) or the corals for homologous Symbiodiniaceae isolation (HOM) were maintained at.

**Table 2.** Details of the coral groups in the heat stress experiment.

| **Coral group** | **Dominant Symbiodiniaceae species** | **Menthol treatment** | **Coral inoculation** | **Coral genotype** | **Tank n** | **Fragment n** | **Polyp n** |
| --- | --- | --- | --- | --- | --- | --- | --- |
| SS8-corals | *Cladocopium proliferum* C1 | Yes | Yes- heat-evolved, cultured | G3, G4 | 4 | 24 | 312 |
| WT10-corals | *Cladocopium proliferum* C1 | Yes | Yes- wild-type, cultured | G3 | 2 | 14 | 168 |
| HOM-corals | *Durusdinium trenchii* and *D. glynnii* D1 | Yes | Yes- freshly isolated from *G. fascicularis* | G3, G4 | 6 | 36 | 396 |
| Native-corals | *Durusdinium trenchii* and *D. glynnii* D1 | No | No | G3, G4 | 6 | 36 | 306 |
| NativeC-corals | *Cladocopium* C40 | No | No | G2 | N/A* | 10 | 85 |

\* due to the low number of replicates of NativeC-corals, Native-corals and NativeC-corals were in the same treatment tanks. Also note that NativeC-corals were only used for mass spectrometry imaging.

Coral inoculation was carried out in four replicate tanks per Symbiodiniaceae treatment and with nine coral fragments per tank (i.e., 3 Symbiodiniaceae treatments x 4 tanks x 9 coral fragments = 108 total) (Fig. 1). Three inoculations were conducted over the course of seven weeks (week 0, 4 and 7; Fig. 1, Table S1). On the day of inoculation, the circulating pumps were stopped, and the water volume of the tanks was reduced to 4 L to allow high densities of the algal inocula. The inocula were added to yield a final density of ∼10^4^ cells mL^-1^; Shellfish Diet 1800® was added at 5000 cell mL^-1^ to encourage phagocytosis. After 5 h, the water volume was increased to 8 L and the circulating pumps were restarted. The normal water volume (13.5 L) was resumed the following day. Eighteen coral fragments were kept as negative control (i.e., chemically bleached but no inoculation). The corals were allowed to recover for 18 weeks after the first inoculation to regain the Symbiodiniaceae population prior to a short-term heat stress experiment (Fig. 1). Due to the relatively low light intensity that the cultured Symbiodiniaceae were exposed to (30-60 µE photons m^-2^ s^-1^), the light intensity of the aquaria was reduced to 45-55 µE m^-2^ s^-1^ at full sun until the completion of the last inoculation; thereafter light levels were gradually increased to 80-85 µE m^-2^ s^-1^ at full sun (Table S3). Corals were fed three times a week and 50% of the seawater was changed weekly.

#### 2.1.4 Assessment of inoculation outcomes via DNA metabarcoding

Four sets of coral samples, each containing one polyp, were taken to verify the inoculation outcomes via metabarcoding. These included 1) each coral genotype shortly after collection from the field (1 replicate x 5 genotypes, *n* = 5), 2) corals at the end of menthol treatment (*n* = 4), 3) coral at seven weeks after the first inoculation (4 replicates x 4 tanks x 4 coral groups + 8 negative control, *n* = 72), and 4) corals at 18 weeks after the first inoculation (6 replicates x 4 coral groups, *n* = 24) (Table S1). In addition, 1 mL of each inoculum was sampled (10^4^ cell mL^-1^, 3 replicates x 3 inocula, *n* = 9) and three 50:50 *Cladocopium*: *Durusdinium* mock communities were made. (Supplementary Methods S2). See Supplementary Methods S3 for DNA extraction, PCR amplification and library preparation procedures. Samples were sequenced at the Walter and Eliza Hall Institute with Illumina MiSeq v3 and raw sequences were submitted to SymPortal (Hume et al. 2019) for Symbiodiniaceae community analysis.

### 2.2 Heat stress experiment

#### 2.2.1 Experimental design

The design of the heat stress experiment was based on the results of the Symbiodiniaceae community analysis post-inoculation (see Results-Inoculation outcome) and included five coral groups: 1) SS8-corals, 2) WT10-corals, 3) HOM-corals, 4) Native-corals and 5) NativeC-corals (Fig. 1; Table 2). A total of 6-7 coral fragments were randomly and evenly distributed into each of the ambient (27°C) and elevated (32°C) treatment tank (replicate tank *n* = 4 for SS8-corals, *n* = 2 for WT10-corals, *n* = 6 for HOM-corals, *n* = 6 for Native-corals and NativeC-corals) (Table 2). WT10-corals had lower replication since only one coral genotype (G3) acquired this inoculum. NativeC-corals were only used for metabolite profiling as only 10 fragments total were available. For the elevated treatment, the temperature was ramped up 1°C day^-1^ until reaching 32°C, which was then maintained for 8 days. Since corals may exchange Symbiodiniaceae when in close proximity with each other, corals of different Symbiodiniaceae treatments were maintained in their respective tanks. The seawater temperature and carbonate chemistry conditions are shown in Table S4.

#### 2.2.2 Symbiodiniaceae community and physiological measurements

Sampling for Symbiodiniaceae community, Symbiodiniaceae cell density, photochemical efficiency, coral survival and size was conducted at the beginning (i.e., before temperature ramping, at week 18 after the first inoculation) and the end (i.e., after eight days under 32°C, at week 20) of the experiment (Fig. 1, Table S1). One coral polyp each was sampled for Symbiodiniaceae community analysis (6 replicates x 4 coral groups x 2 temperatures, *n* = 48) and cell density measurement (5 replicates x 4 coral groups x 2 temperatures, *n* = 40). Because SS8 relative abundance in corals observed at week 18 was highly variable, six extra samples were collected under elevated temperatures. Cell count, DNA extraction, PCR amplification and library preparation were conducted as before. Coral size and survival were determined by polyp count using all corals (SS8-corals *n* = 24, WT10-corals *n* = 14, HOM-corals *n* = 36, Native-corals *n* = 36). A polyp was counted as ‘dead’ when > 50 % of its tissue was lost, and a new grown polyp was counted as ‘alive’ when its six primary septa were fully formed. Photochemical efficiency (dark-adapted or maximum quantum yield of photosystem II; Fv/Fm) of all corals was measured using the Walz Imaging Pulse Amplitude Modulation (I-PAM) Fluorometer driven by the software ImagingWin. The corals were dark-adapted for 20 min prior to measurement; three technical replicate measurements were taken per coral fragment and measurements were always conducted at 09:00 am. Additional measurements for photochemical efficiency were taken 4 days after exposure to 32°C.

### 2.3 Statistical analysis

#### 2.3.1 Symbiodiniaceae community and cell count

Raw sequences were submitted to SymPortal (Hume et al. 2019) for Symbiodiniaceae ITS2 taxa and profile analysis. Data visualization and statistical analysis was conducted in R (version 4.1.2) with a customed pipeline. Samples with low reads (four samples, < 500 reads each) were excluded from the analysis. PERMANOVA (Oksanen et al. 2016) based on unweighted UniFrac distance and 999 permutations was applied to compare the Symbiodiniaceae communities of the coral groups between ambient and elevated temperatures. Multivariate homogeneity of variances was tested and confirmed using *betadisper* (Anderson et al. 2006). The effect of coral group and temperature treatment on Symbiodiniaceae cell density was analyzed using negative binomial generalized linear models and model fitness was confirmed with chi-square test (Bolker et al. 2009).

#### 2.3.2 Coral growth, survival and photochemical efficiency

Since all coral fragments had 3-4 polyps at the time of inoculation (week 0), the number of polyps at week 18 (coral size) can be directly used for comparison, with no normalization required. The size data was normally distributed (Shapiro test) and with equal variance (Levene’s test); the effect of coral group was tested with an ANOVA followed by TukeyHSD for pairwise comparison. Coral survival was calculated based on the increase or decrease in the number of polyps (i.e., number of polyps at the end of the experiment minus the starting number of polyps). The effect of coral group and temperature treatment was analyzed separately using the non-parametric Kruskal-Wallis test, followed by Dunn’s test (Dunn 1964) for pairwise comparisons. The p-values were corrected using the Benjamini–Hochberg method (Benjamini and Hochberg 1995). The percentage of change was used for visualization (i.e., number of polyp change divided by the starting number of the polyps). For photochemical efficiency, the three technical replicate measurements of each sample were first merged before statistical analysis. Beta regressions were applied to test the effect of the interaction of temperature and coral group on photochemical efficiency on day 8 and day 4 since exposure to 32°C using the package *betareg* (Cribari-Neto and Zeileis 2010). The models were inspected with the *plot* function and met the model assumptions.

### 2.4 High-resolution mass spectrometry imaging

#### 2.4.1 Sample collection and fourier transform ion cyclotron resonance (FT-ICR) matrix-assisted laser desorption/ionization mass spectrometry imaging (MALDI-MSI)

Three samples of each coral group, each with two technical replicates, were collected for metabolite profiling with MSI after 8 days under 27 and 32°C (3 biological replicates x 2 technical replicates x 5 coral groups x 2 temperatures = 60 total). To better identify host metabolites, 3 samples of chemically bleached corals were collected immediately after four weeks of menthol treatment. Each sample consisted of one coral polyp that was removed with a bone cutter. Sample preparation, cryosectioning, matrix spray and mass spectrometry imaging were conducted following the methods in (Chan et al. 2023) (Supplementary Methods S4). Briefly, coral samples were anaesthetized with MgCl_2_, quickly rinsed with MQ water, embedded in a carboxymethyl cellulose (CMC) and gelatin mixture, snap frozen on dry ice and stored at −80°C (Fig. 2). The samples were cryosectioned (Leica CM 1860, −25°C) until a substantial amount of the frozen body and tentacles were exposed. A thin layer (∼2 µm) of the CMC and gelatin mix was applied on the exposed sample to strengthen its integrity before collecting a section. Four consecutive sections were collected per sample on a cryofilm at 12 µm thickness, freeze-dried, mounted on a stainless-steel sheet, and sprayed with α-Cyano-4-hydroxycinnamic acid (HCCA). MALDI-MSI analysis was conducted on the Bruker SolariX (7T XR hybrid ESI–MALDI–FT–ICR–MS) (150-2000 m/z, 50 µm spatial resolution, positive ion mode). SCiLS was used for noise removal and peak picking, and the final peak list consists of 418 metabolites. Their intensities (average peak area) were normalized with root mean square and with a section’s total surface area.

**Fig. 2.**
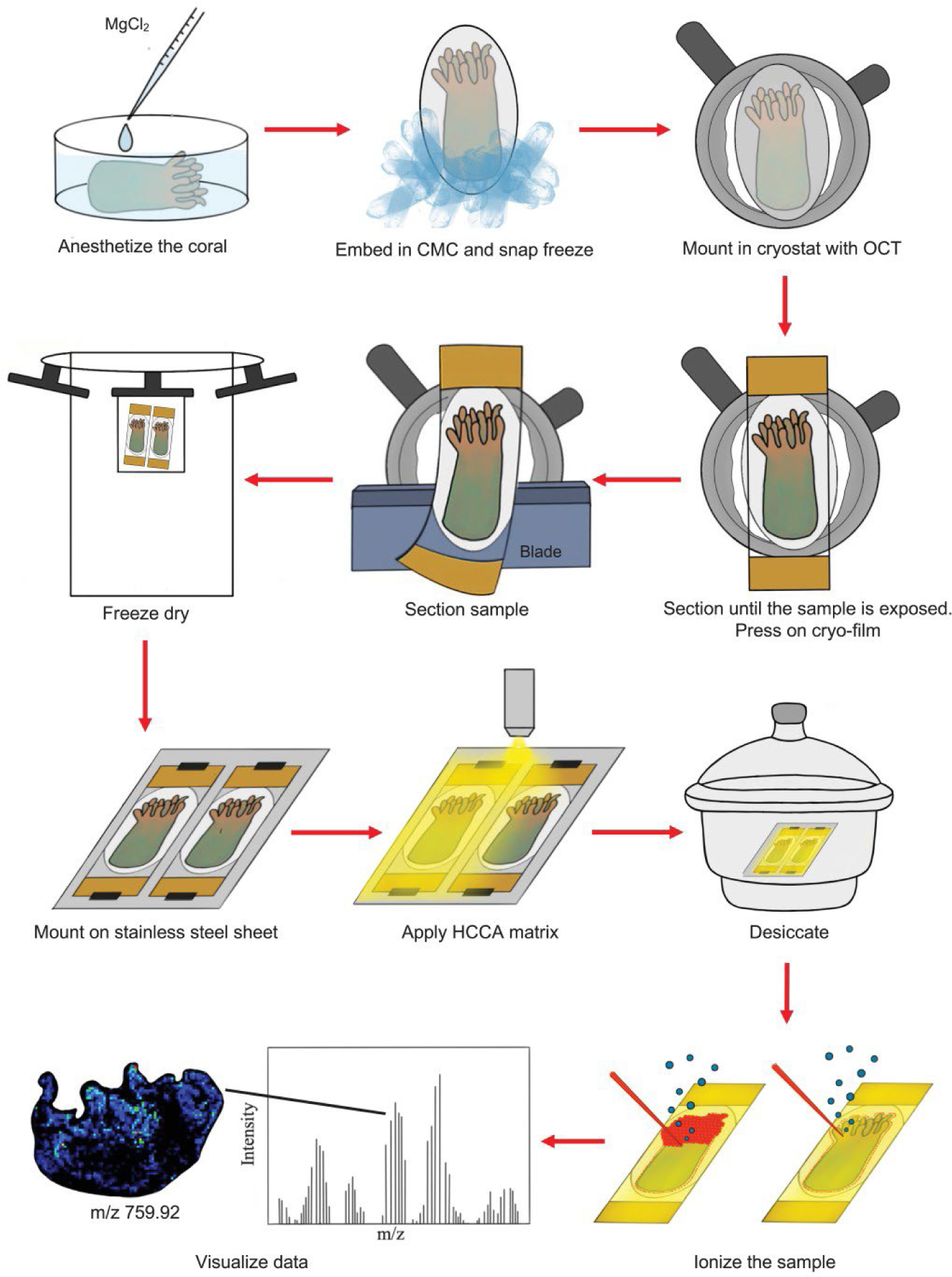
Fourier transform ion cyclotron resonance (FT-ICR) matrix-assisted laser desorption/ionization mass spectrometry imaging (MALDI-MSI) workflow of the coral samples. CMC = 2% carboxymethyl cellulose with gelatin; OCT = optimal cutting temperature compound; HCCA = α-Cyano-4-hydroxycinnamic acid.

#### 2.4.2 Metabolite annotation

Metabolites were annotated with 1) an in-house library, established with MALDI-thin layer chromatography on Symbiodiniaceae and the coral model sea anemone (Chan et al. 2023); 2) HPLC-UV profiling of natural pigments with absorption maxima at 430 and 662 nm; 3) concurrent liquid-chromatography mass spectrometry analysis on *G. fascicularis*; and 4) annotation against the LipidMaps, ChEBI and HMDB databases within METASPACE, with reference to known coral-symbiont metabolites (Roach et al. 2021). For pigment annotations, three sets of samples, each containing four coral polyps were collected, covering WT10-corals, HOM-corals and chemically bleached corals. Note that these samples were used for pigment annotation only and were not intended for quantitative comparison between experimental groups.

#### 2.4.3 Statistical analysis

The normalized data was analyzed in MetaboAnalyst 5.0 with log transformation and no data scaling, and data normality and homogeneity confirmed visually. Firstly, the data was divided into ambient and elevated temperatures to examine the effect on coral groups (SS8-, WT10-, HOM-, Native-, NativeC-corals) within a temperature treatment. PCAs were generated using all 418 metabolites detected, and the effect of coral groups was tested by a one-way ANOVA. The top 60 most significant metabolites (in terms of differences in their relative intensities between experimental groups) from the ANOVA were then shown on a heatmap, with Euclidean distance and Ward clustering algorithm applied. Next, the effect of temperature (27°C, 32°C) within a coral group was examined using t-tests. A metabolite was considered significant when P_adj_ (Benjamini and Hochberg 1995) was < 0.05 and fold change (FC; calculated based on the non-log transformed data) was > 1.3 (i.e., > 30% difference). All significant metabolites detected were then visualized on a heatmap using Euclidean distance and Ward clustering algorithms. A PCA was generated using all 418 metabolites, for comparison with the heatmaps that only showed significant metabolites. The localization and relative intensities of metabolites with significant differences were then visualized on SCiLS.

### 2.5 Long-term monitoring

To track the long-term stability of symbiosis with SS8, SS8-corals were kept under ambient temperature (27°C) and sampled at one year (*n* = 4) and two years (*n* = 4) after the first inoculation, for Symbiodiniaceae community analysis. WT10-coral had no survivors by year one.

## 3. RESULTS

### 3.1 Coral inoculation experiment

#### 3.1.1 Efficacy of menthol treatment

Four weeks of menthol treatment removed ∼98.8% of *G. fascicularis*’ native Symbiodiniaceae. On average, control corals (non-menthol-treated) contained ∼2,730,000 Symbiodiniaceae cells per polyp, whereas menthol-treated corals had ∼33,000 cells per polyp (i.e., 121 times fewer cells compared with the control) (Fig. 3). Coral genotype had a significant effect on Symbiodiniaceae cell density in control corals (Kruskal-Wallis, χ^2^ = 10.79, df = 3, *p* = 0.013) and menthol-treated corals (Kruskal-Wallis, χ^2^ = 12.49, df = 3, *p* = 0.005). In both cases, genotypes G3, G4 and G5 had similar cell densities while G1 had a higher cell density than G3 and G4 (Fig. 3, Table S5). Menthol treatment removed 97.8% of G1’s Symbiodiniaceae population, but removed 99.1-99.2% of G3, G4 and G5’s Symbiodiniaceae population. Menthol treatment resulted in little coral mortality (0-4.2%), except in one coral genotype (G2, 60% mortality), which was excluded from downstream physiological measurements due to a limited number of replicates remaining (Table S2).

**Fig. 3.**
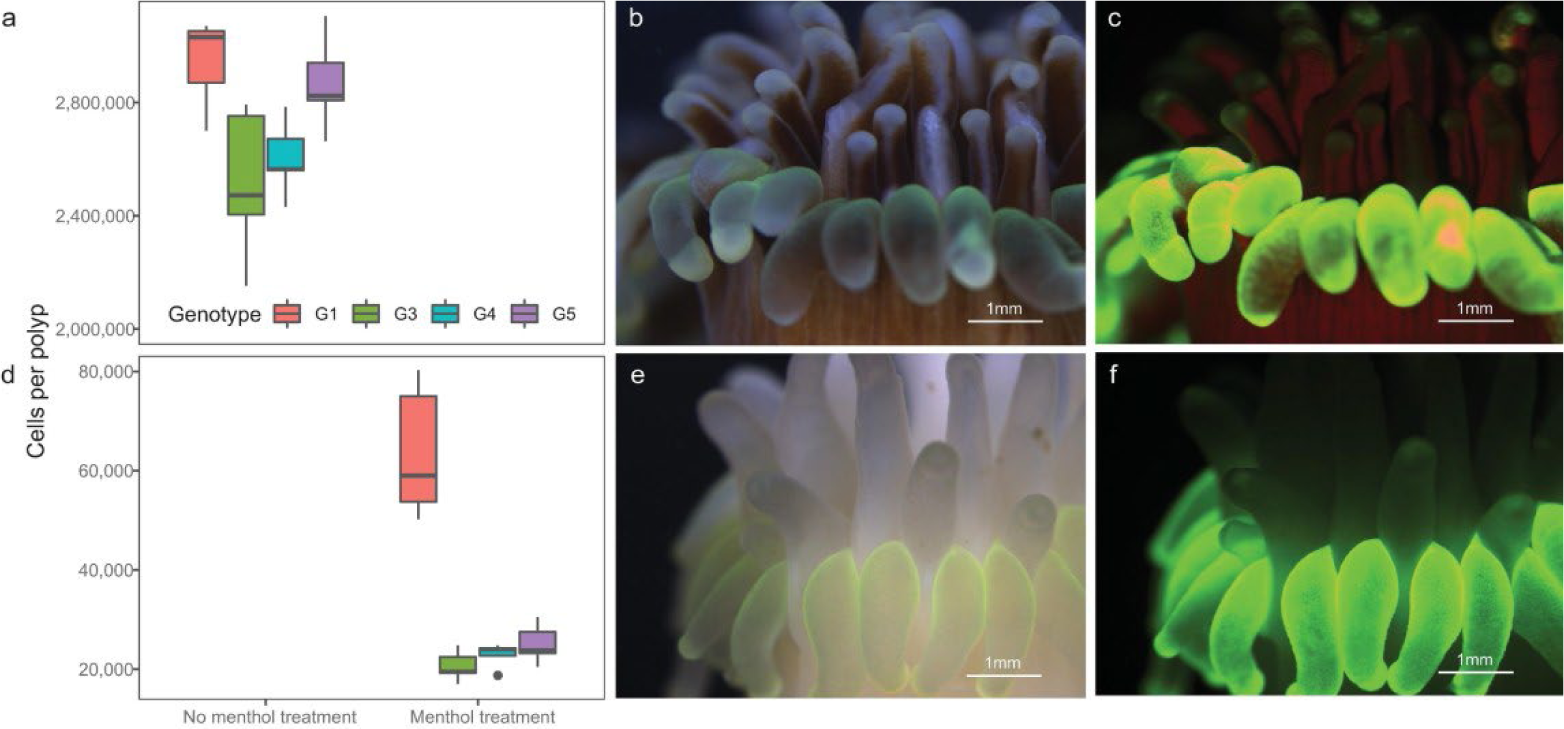
Symbiodiniaceae cell densities in control and menthol-treated fragments of *Galaxea fascicularis*. a) cell density, b) bright field image and c) fluorescent image of a control *G. fascicularis* polyp (non-menthol-treated). d) cell density, e) bright field image and f) fluorescent image of a *G. fascicularis* polyp after four weeks of menthol treatment. The images represent coral genotype G3 and were taken with a Leica M205 FA using a green fluorescence protein filter.

#### 3.1.2 Inoculation outcome

ITS2 metabarcoding was used to assess the Symbiodiniaceae community composition in experimental coral fragments. The average read depth across all samples was ∼16,000. No ITS2 profile was identified in any DNA extraction and PCR negative control (average < 55 reads), indicating there was no contamination. The native Symbiodiniaceae community of *G. fascicularis* was dominated by *D. trenchii* and *D. glynnii* (87-97%; ITS2 profile: D1-D4-D4c-D1h-D1c, D1-D4-D4c-D1c-D1cj, D1-D4-D4c-D6-D1c-D2, or D1-D4-D2-D4c-D6), with the exception of genotype G2 being dominated by *Cladocopium* C40 (72%; ITS2 profile: C40-C3-C115-C40h) (Fig. S1). These native Symbiodiniaceae populations are hereafter referred to as “native *Durusdinium*” and “native *Cladocopium*”. The ITS2 profile of the inocula SS8 and WT10 (C1-C1b-C1c-C42.2-C1bh-C1br-C1cb-C3) was distinct from the native Symbiodiniaceae communities of *G. fascicularis*, therefore detection of this profile is a reliable indicator of successful inoculation (Fig. 4). The freshly isolated Symbiodiniaceae from *G. fascicularis* (HOM inoculum) contained mixed native *Durusdinium* (75-85%) and native *Cladocopium* (9-18%) (Fig. 4).

**Fig. 4.**
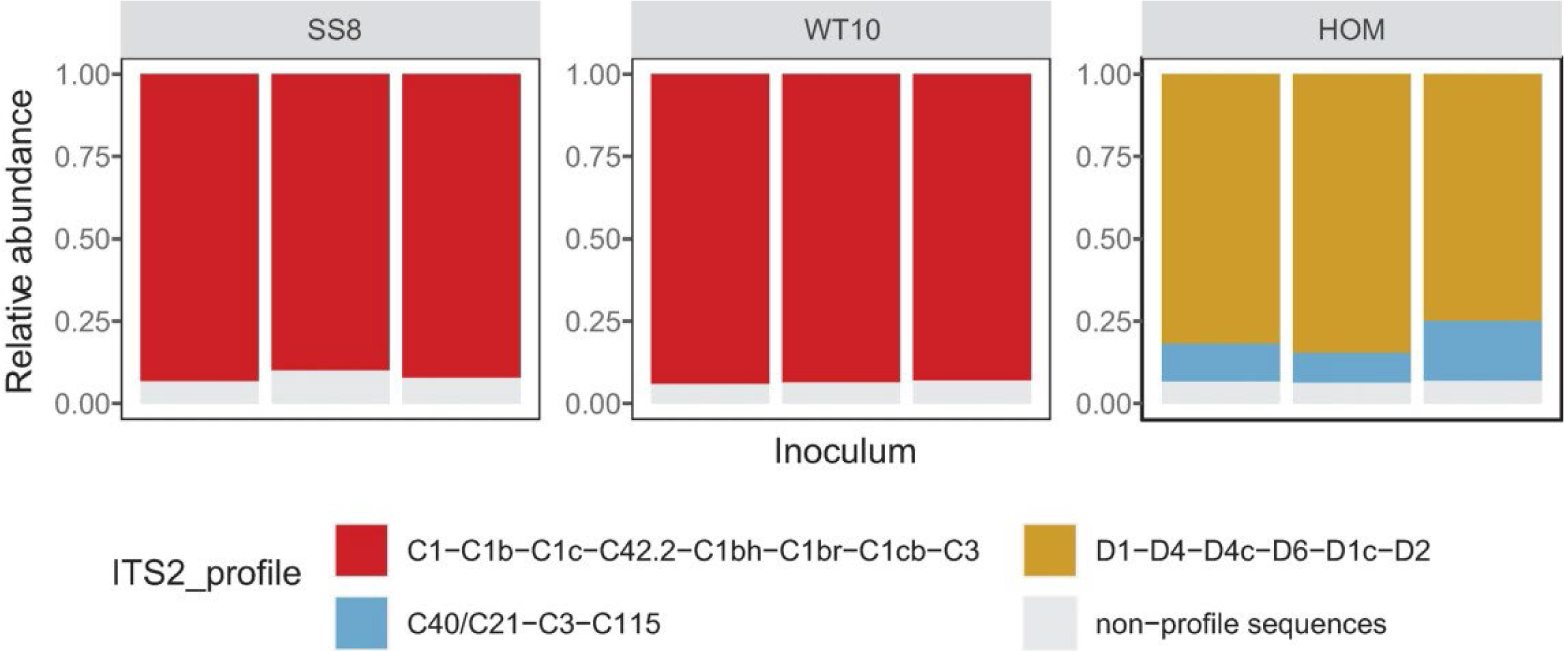
ITS2 profile of the inocula. SS8 = heat-evolved *Cladocopium proliferum*; WT10 = wild-type *C. proliferum*; HOM = homologous inoculum freshly isolated from *G. fascicularis*.

Seven weeks after the first inoculation, two out of the four *G. fascicularis* genotypes (G3 and G4) contained detectable levels of SS8 or WT10 (Fig. S2), hence only these genotypes were retained for the experiment. The proportion of SS8 or WT10 accounted for ∼1-15% of the corals’ Symbiodiniaceae communities. By 18 weeks after the first inoculation, SS8 accounted for an average of 56% (ranged 4-87%) in SS8-corals, and WT10 accounted for an average of 69% (ranged 54-81%) in WT10-corals (Fig. 5). HOM-corals had a similar Symbiodiniaceae community to the Native-corals (which was not bleached nor inoculated), both being dominated by native *Durusdinium*. *Cladocopium* C40 had minimum relative abundance in HOM-corals, suggesting the C40 component of the HOM inoculum (9-18%) had no or little infection success.

**Fig. 5.**
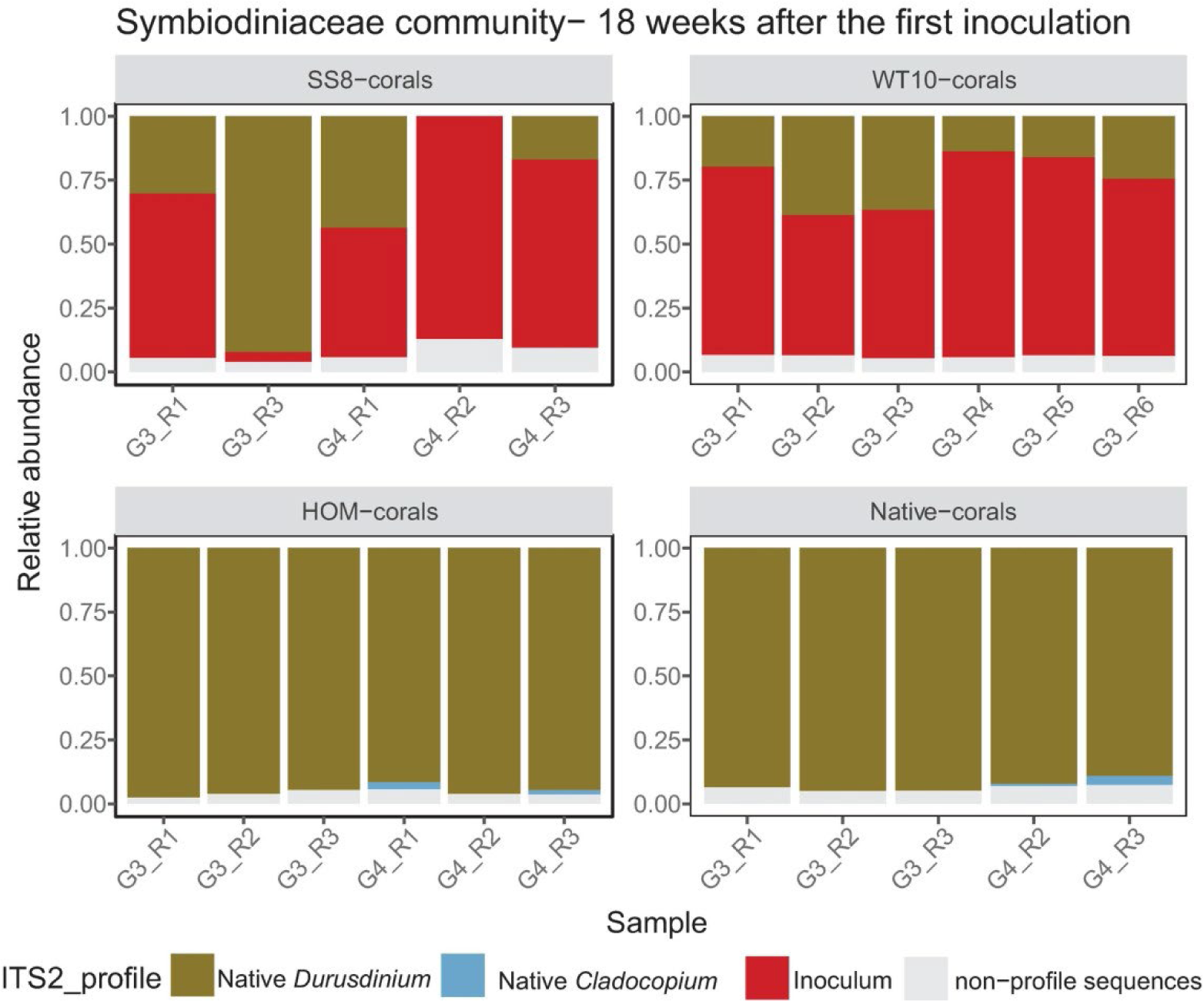
Coral-associated Symbiodiniaceae communities 18 weeks after the first inoculation. The Symbiodiniaceae treatments of the corals included: heat-evolved *Cladocopium proliferum* (SS8-corals), wild-type *C. proliferum* (WT10-corals), freshly isolated, homologous Symbiodiniaceae from *Galaxea fascicularis* (HOM-corals). Control corals with no menthol treatment nor inoculation are also included (Native-corals). All corals were grown under ambient temperature during this period. “G” refers to the sample’s genotype, “R refers to the sample’s replicate number. Note that one SS8- and Native-coral sample were excluded due to low read numbers.

### 3.2 Heat Stress Experiment

#### 3.2.1 Symbiodiniaceae community

At the end of the heat stress experiment (i.e., 8 days under 32°C), the Symbiodiniaceae communities of WT10-corals under ambient and elevated temperatures were significantly different (PERMANOVA, *p* = 0.004). The proportion of WT10 in coral fragments at ambient temperature (79%) was on average 21% higher compared to that at elevated temperature (58%) (Fig. 6). In contrast, the Symbiodiniaceae communities of SS8-, HOM- and Native-corals remained unchanged under the two temperatures (PERMANOVA, *p* = 0.50, 0.83, 0.97 respectively; Fig. 6). In SS8-corals, a high level of variability in SS8 relative abundance (ambient: 43-80%; elevated: 24-73%) was observed, whereas limited variability was found in WT10-corals (ambient: 73-82%; elevated: 50-64%). The Symbiodiniaceae community of HOM- and Native-corals comprised of nearly 100% native *Durusdinium* under both temperatures.

**Fig. 6.**
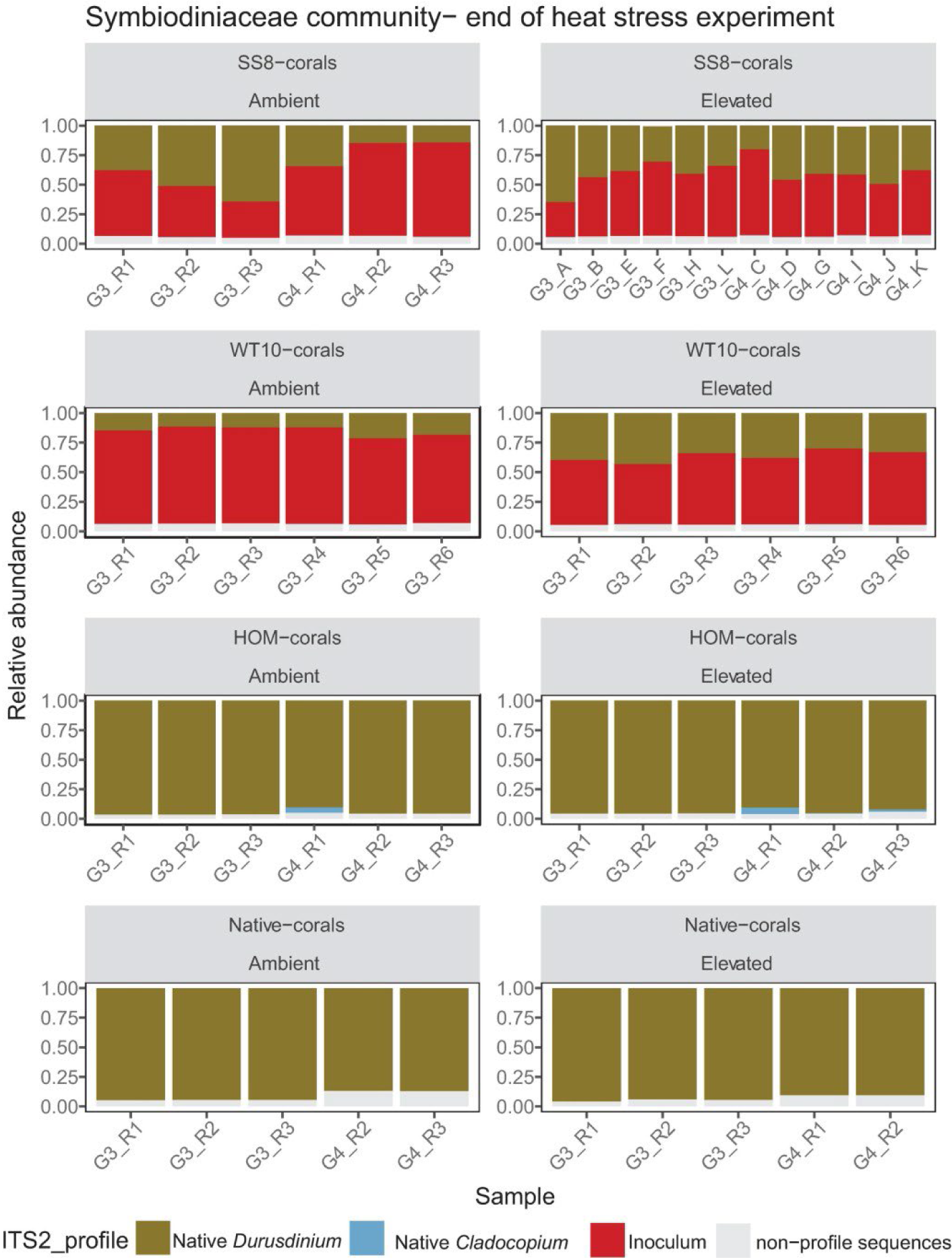
Coral-associated Symbiodiniaceae communities at the end of the short-term heat stress experiment (8 days, 32°C), 20 weeks after the first inoculation. The abbreviations of the experimental groups refer to corals that were inoculated with and dominated by heat-evolved *Cladocopium proliferum* (SS8-corals), wild-type *C. proliferum* (WT10-corals), freshly isolated, homologous Symbiodiniaceae from *Galaxea fascicularis* (HOM-corals); as well as control corals with no menthol treatment nor inoculation (Native-corals). “G” refers to the sample’s genotype, “R refers to the sample’s replicate number. An extra six samples were collected for SS8-corals under elevated temperature due to high level of variability in SS8 relative abundance observed in week 18 (Fig. 5), with replicate numbers indicated as A-K. Note that one Native-coral sample under ambient and elevated temperatures was excluded due to low read number.

#### 3.2.2 Symbiodiniaceae cell density

Under elevated temperature, WT10-corals had ∼89% lower Symbiodiniaceae cell density when compared to their ambient counterparts (GLM, *z* = −39.14, *p* < 0.001). Cell density in this group was the lowest of all experimental groups (Fig. 7a; Table S6). Native-corals also had a ∼11% decrease in cell density at elevated compared with ambient temperature (GLM, *z* = −2.13, *p* = 0.033), whereas SS8- and HOM-corals showed no difference in cell density between temperatures (GLM, *p* = 0.95, 0.60 respectively) (Fig. 7a). Under elevated temperature, the cell density of SS8-corals was similar to that of Native-corals which did not undergo menthol treatment (GLM, *z* = 1.38, *p* = 0.167). Under ambient temperature, all coral groups that underwent menthol treatment and inoculation (SS8-, WT10-, HOM-corals) had lower cell densities than Native-corals (GLM, *p* < 0.001 for all, Table S6), suggesting they had not fully recovered from chemical bleaching. By 20 weeks after the first inoculation, both SS8- and WT10-corals had recovered to ∼82% of their original cell density prior to menthol treatment, while HOM-corals only recovered to ∼65% (Fig. 7b), despite these groups having been inoculated at the same cell density throughout the three inoculations. Cell densities of SS8- and WT10-corals were higher than those of HOM-corals under ambient temperature (Fig. 7a; Table S6).

**Fig. 7.**
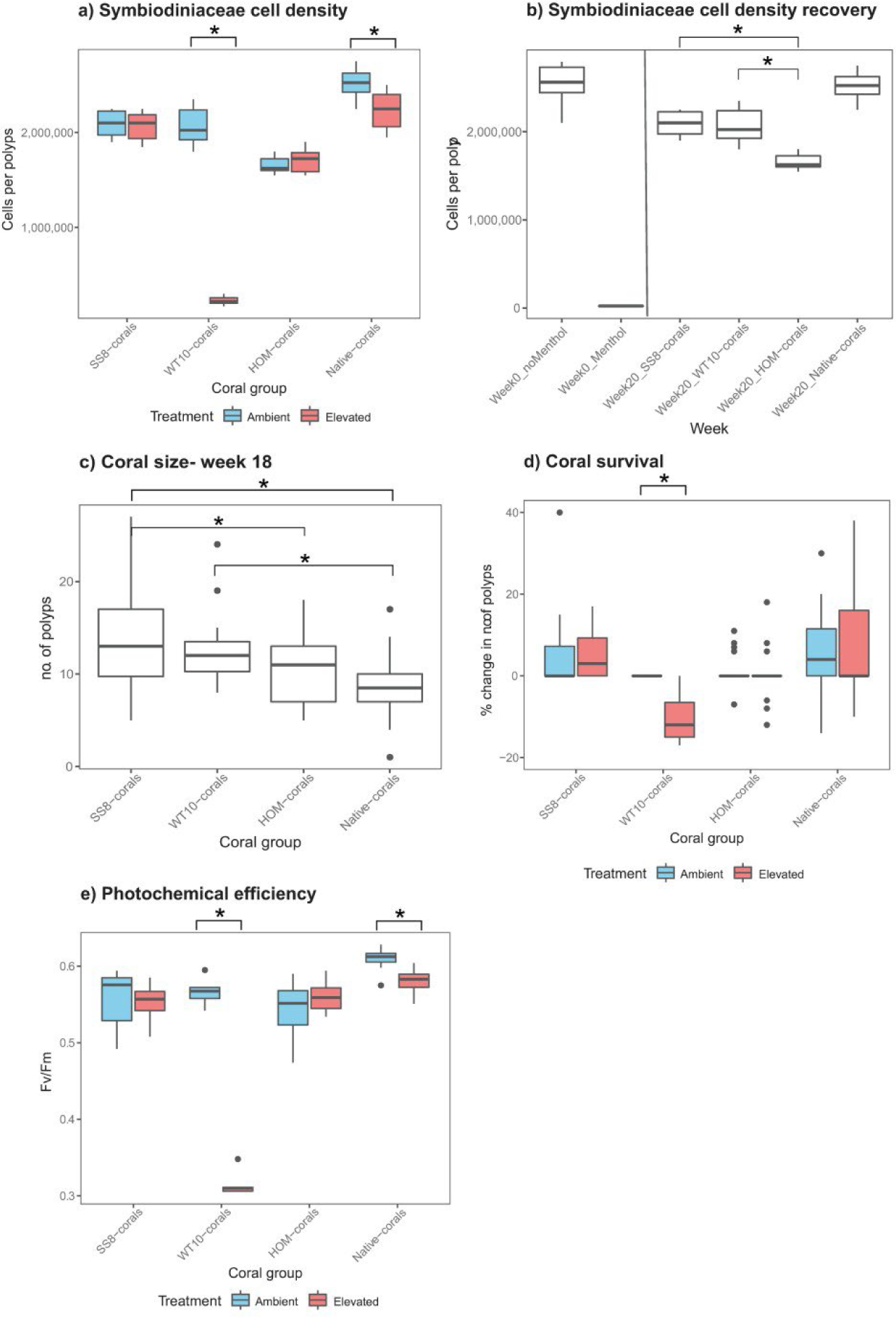
Physiological responses and coral size of the four coral groups. a) Symbiodiniaceae cell density per coral polyp under ambient and elevated temperatures. b) Symbiodiniaceae cell density per coral polyp before, immediately after, and 20 weeks after menthol treatment (at the end of the heat stress experiment). c) coral size (number of polyps) 18 weeks after first the inoculation; corals were exposed to ambient temperatures during this time. d) coral survival, presented as the percentage of change in the number of polyps. e) coral photochemical efficiency (maximum quantum yield, Fv/Fm). The Symbiodiniaceae treatment of the coral group included: heat-evolved *Cladocopium proliferum* (SS8-corals), wild-type *C. proliferum* (WT10-corals), freshly isolated, homologous Symbiodiniaceae from *Galaxea fascicularis* (HOM-corals). Control corals with no menthol treatment nor inoculation that were natively dominated by Durusdinium sp. were also included (Native-corals). * indicates statistical significance. For simplicity, statistical significance is only shown for temperature effect in a), d), e) and not for coral group effect.

#### 3.2.3 Coral size and survival

All coral fragments had 3-4 polyps at the time of the first inoculation. They were grown under ambient temperature for an 18-week recovery period before the heat stress experiment. Over the recovery period, coral groups differed significantly in growth rate as assessed by their size at the end of this period (i.e., the total number of polyps per coral fragment) (ANOVA, F = 8.57, *p* < 0.001). Both SS8- and WT10-corals grew more than Native-corals (TukeyHSD, *p* < 0.001, 0.009); SS8-corals also grew faster than HOM-corals (TukeyHSD, *p* = 0.017) (Fig. 7c; Table S7). The median size after the 18-week period was 13 polyps (SS8-corals), 12 polyps (WT10-corals), 11 polys (HOM-corals) and 8.5 polyps (Native-corals) (Fig. 7c). Survival over this period was 100% for all coral groups. The change in size of coral fragments over the 10-day period of exposure to elevated temperature was too small to be reliably measured with the tools that were available for this experiment.

Survival over the course of the heat stress period did not differ with survival in the ambient control for all groups, except for WT10-corals which had significantly higher mortality (12%) compared to its ambient counterparts (Kruskal-Wallis, χ^2^ = 9.02, df = 1, *p* = 0.003). Under elevated temperature, there was a significant effect of coral group on survival (Kruskal-Wallis, χ^2^ = 20.07, df = 3, *p* < 0.001), where SS8-, HOM- and Native-corals survived better than WT10-corals (Dunn test, *p* < 0.001, 0.015, < 0.001 respectively) (Fig. 7d; Table S8).

#### 3.2.4 Photochemical efficiency

Under elevated temperature, photochemical efficiency of SS8- and HOM-corals did not differ from their ambient counterparts (beta regression, *p* = 0.56, 0.06, respectively) (Fig. 7e; Table S9). In contrast, both WT10- and Native-corals had lower photochemical efficiency under elevated temperatures than under ambient temperatures (beta regression, *p* < 0.001 for both) (Fig. 7e; Table S9). This pattern was already significant halfway through the heat stress experiment (day 4) (Fig. S3). At the end of the experiment, WT10-corals exhibited a severe 44% difference in photochemical efficiency between ambient (Fv/Fm averaged 0.57) and elevated temperature (0.32), while Native-corals showed a minor 5% difference (ambient averaged 0.61; elevated 0.58). Under elevated temperature, SS8-, HOM- and Native-corals had higher photochemical efficiency than WT10-corals (beta regression, *p* < 0.001 for all, Table S9). Under ambient temperature, the photochemical efficiency of HOM-corals was slightly lower than that of WT10-corals (beta regression, *p* = 0.047, difference = 0.02).

The history of menthol treatment had a small yet significant effect on coral photochemical efficiency under ambient temperature (beta regression, *p* < 0.001 for all, Fv/Fm difference = 0.06) (Table S9). Despite the drop in photochemical efficiency in Native-corals under elevated temperature, Native-corals had higher photochemical efficiency than all coral groups (SS8-, WT10-, HOM-corals) that were treated with menthol and inoculated with Symbiodiniaceae, under both ambient and elevated temperatures (Table S9).

#### 3.2.5 Metabolite profiles

Using mass spectrometry imaging, a total of 418 metabolites were detected across samples following noise removal, 225 (61%) of which were annotated (Table S10). Under elevated temperature, ANOVA detected 182 metabolites with significant differences in relative intensities between SS8-, WT10- and HOM-corals (Table S11). A heatmap of the top 60 significant metabolites showed that half of the SS8-corals replicates clustered with HOM-corals, while the other half clustered with WT10-corals (Fig. S4). T-test showed that only 13 metabolites were significantly different between SS8- and WT10-corals under elevated temperature, with the majority (∼40%) of these being algal pigments localized within the algal symbionts (Figs. 8, 9, S5, Table S12). When these algal pigments were visualized, WT10- and SS8-corals displayed contrasting pigment profiles under elevated temperature, with SS8-corals having a similar profile to HOM-corals (Fig. 8c). One of these pigments (m/z 524.17) was completely absent in WT10-corals under elevated temperature, but was highly abundant in SS8- and HOM-corals (Fig. 8d).

**Fig. 8.**
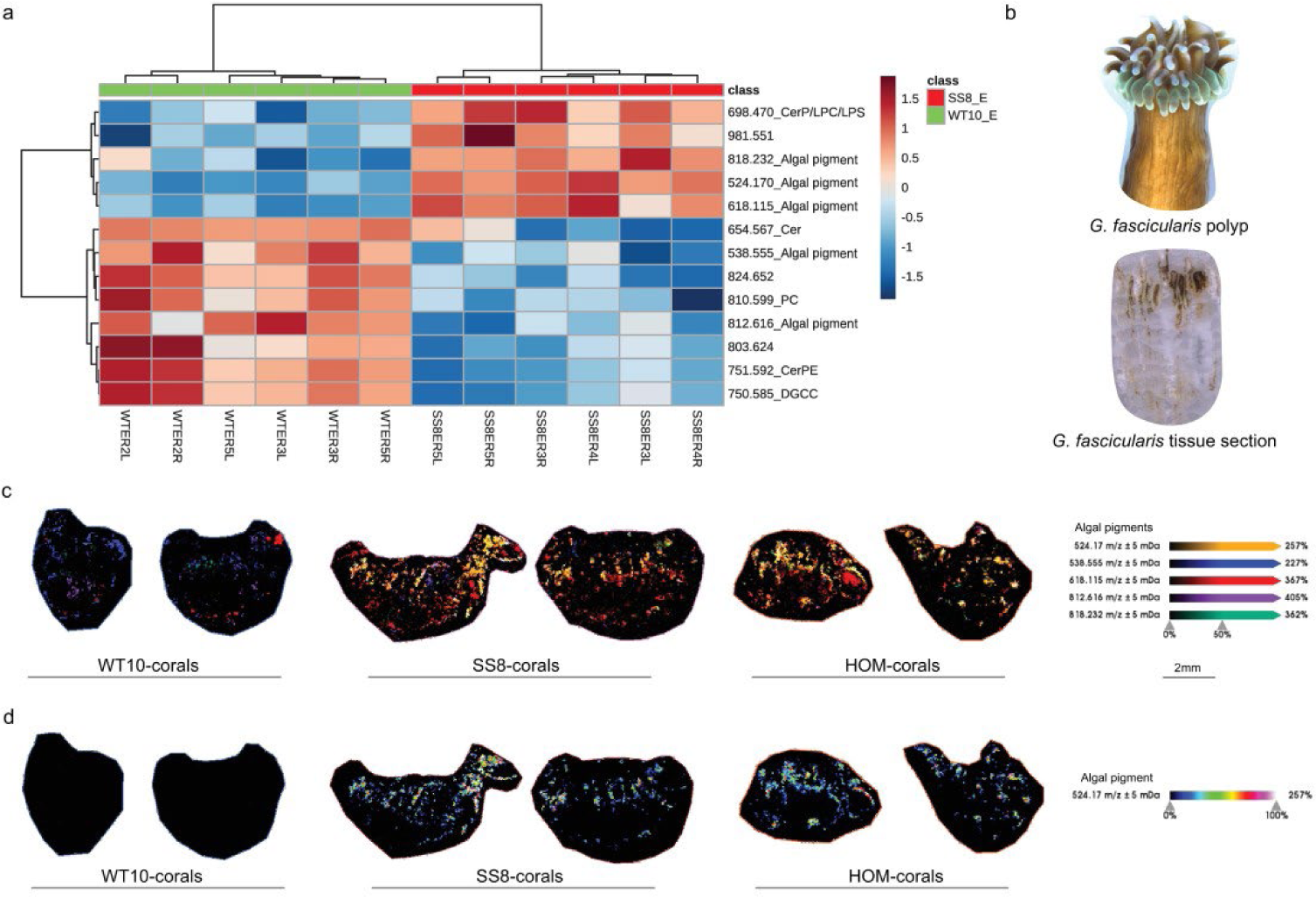
Heatmap and annotations of the 13 metabolites with significant intensity differences between SS8- and WT10-corals under elevated temperature. The color scale indicates log2 fold change relative to the mean. b) images of a *Galaxea fascicularis* polyp and its tissue section at 12 µm thickness. c) algal pigment profile of WT10-corals (hosting wild-type *Cladocopium proliferum*), SS8-corals (hosting heat-evolved *C. proliferum*) and HOM-corals (hosting *Durusdinium trenchii* and *D. glynnii*) under elevated temperatures. d) localization and intensity of the algal pigment m/z 524.17. E = elevated temperature.

**Fig. 9.**
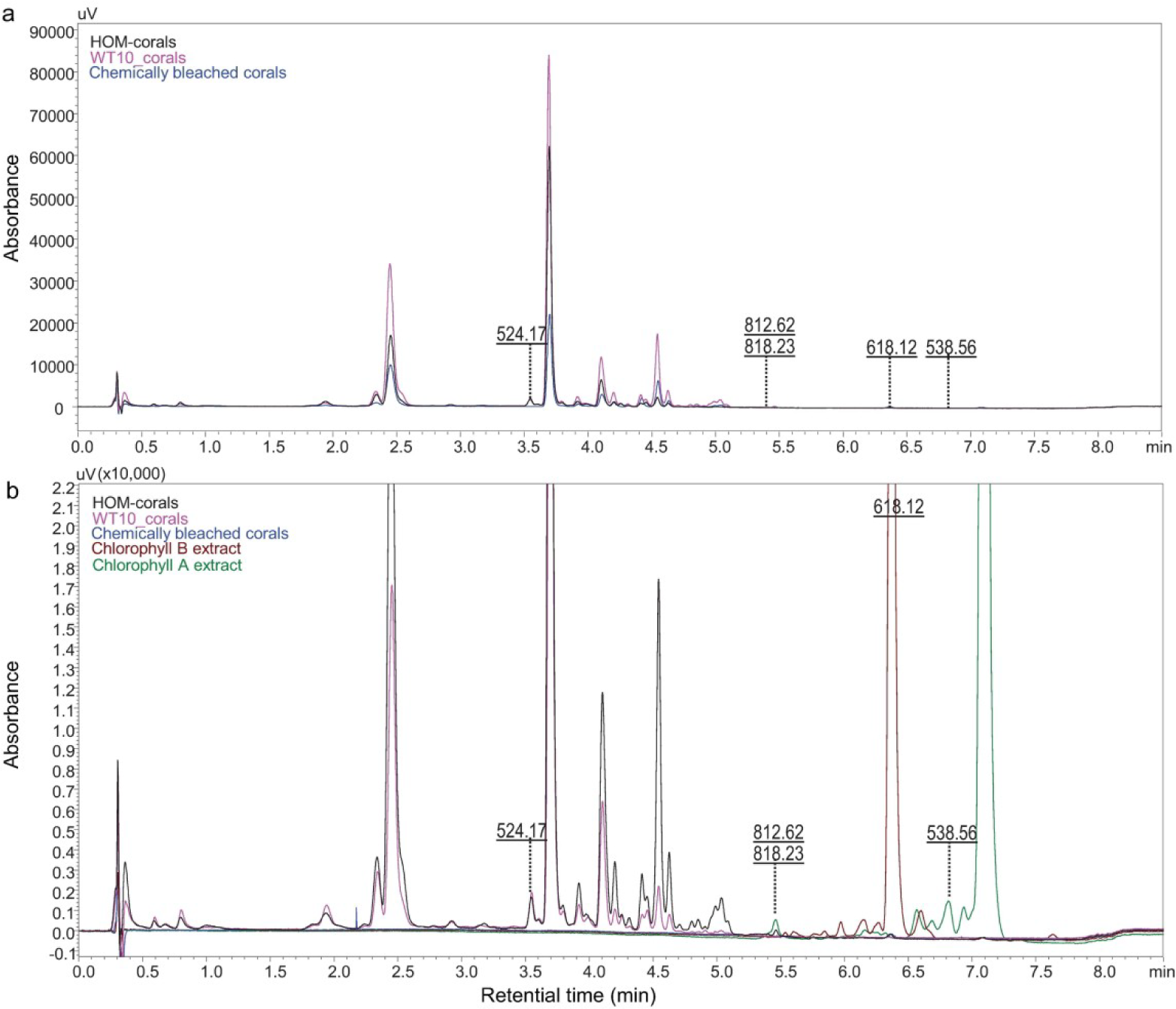
a) HPLC-UV chromatogram absorbing at 430 nm of the lipophilic pigment extracts of three coral samples, run in tandem with LC-MS to identify masses related to pigments. Each sample consisted of four coral polyps pooled from WT10-corals, HOM-corals and chemically bleached corals. Note that these samples were for pigment annotation only and were not intended for quantitative comparison between experimental groups. b) overlay of the pigment profiles of the coral samples (black, pink and blue traces) with chlorophyll B (brown trace) and chlorophyll A (green trace) extracts and their related pigments. The peaks of the algal pigments shown in Fig. 8 are indicated here.

WT10-corals experienced the greatest difference in metabolite profile between ambient and elevated temperatures, with the intensities of 66 out of 418 metabolites (∼16%) being significantly different (Fig. 10a, Table S13). In contrast, SS8-corals only had 12 significantly different metabolites (∼3%) between temperatures (Fig. 10b, Table S14). HOM- and Native-corals, both dominated by *Durusdinium*, showed no difference in metabolite profile between ambient and elevated temperatures. WT10-corals subjected to elevated temperature showed downregulation of three major metabolite classes: glycerophosphocholine (PC, 11 species), ceramide (Cer, 7 species) and betaine lipid (DGCC/MGDG/DGTA/DGTS, 10 species), with a fold change ranging from 1.5 to 81 and with an average of 4.8 (Fig. 10a, Table S13). SS8-corals exposed to elevate temperature also showed downregulation of the same three metabolite classes (PC, 4 species; Cer, 2 species; betaine lipids, 3 species), but the number of significantly different metabolites and their fold change was much smaller (ranging from 1.9 to 4.5 and an average of 3) (Fig. 10b, Table S14). The distributions of PCs overlap completely with coral host tissue and include the boundary of a tissue section, indicating that these are host signals (Fig. 10c). Betaine lipids, in contrast, are concentrated on the coral’s tentacles, with little or no intensity along the tissue section boundary (Fig. 10d). Along the coral tentacles, betaine lipids are found within the location of PCs, but always covering a smaller area than PCs, suggesting that betaine lipids are being packaged inside coral host tissues.

**Fig. 10.**
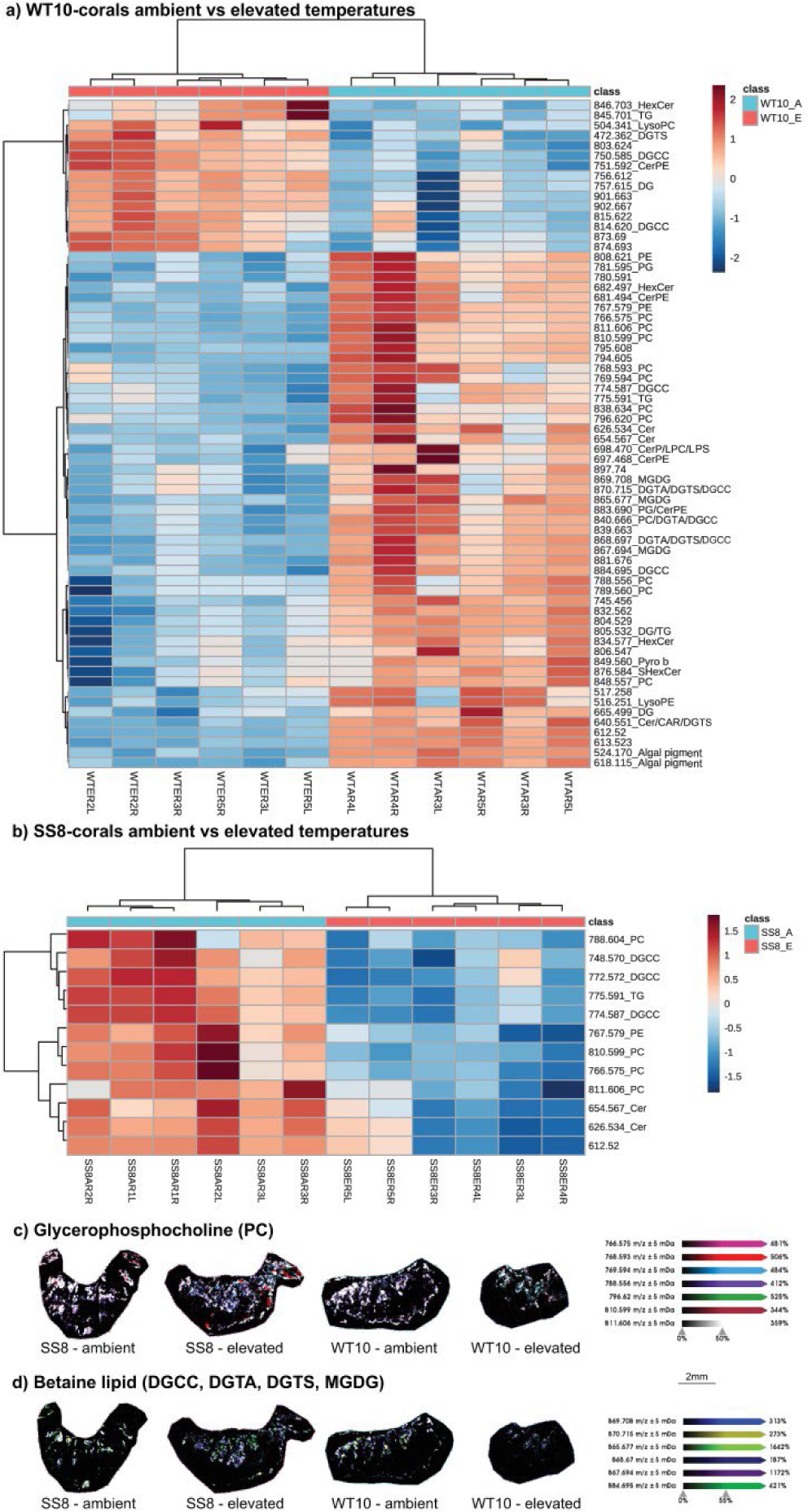
a) Heatmap of the 66 metabolites with significantly different intensities between ambient and elevated temperatures in WT10-corals. b) heatmap of the 12 metabolites with significantly different intensities between ambient and elevated temperatures in SS8-corals. The color scale indicates log2 fold change relative to the mean. A = ambient temperature, E = elevated temperature. c) relative intensities and spatial distribution of glycerophosphocholine (PC) in SS8- and WT10-corals selected from the heatmaps. d) relative intensities and spatial distribution of betaine lipid (DGCC, DGTA, DGTS, MGDG; see Table S10) in SS8- and WT10-corals selected from the heatmaps.

Under ambient temperature, ANOVA detected 308 significant metabolites among coral groups. A heatmap of the top 60 most significant metabolites showed no separation among the menthol-treated/inoculated groups (SS8-, WT10-, HOM-corals), however, clear clustering was observed between this group and control groups (non-menthol-treated; Native-, NativeG2-coral) (Fig. S6). Menthol-treated/inoculated groups consistently showed an upregulation of six ceramides species, compared to the control groups, regardless of their dominant symbiont (*Cladocopium* or *Durusdinium*).

### 3.3 Summary tables

In summary, under ambient temperature WT10- and SS8-corals (both dominated by *C. proliferum*) had similar performance and performed better than HOM-corals (dominated by *D. trenchii* and *D. glynnii*) (Table 3). The highest resilience to elevated temperatures was seen for SS8-corals and HOM-corals, whereas WT10-corals showed signs of stress under elevated temperatures (Table 4).

**Table 3.**
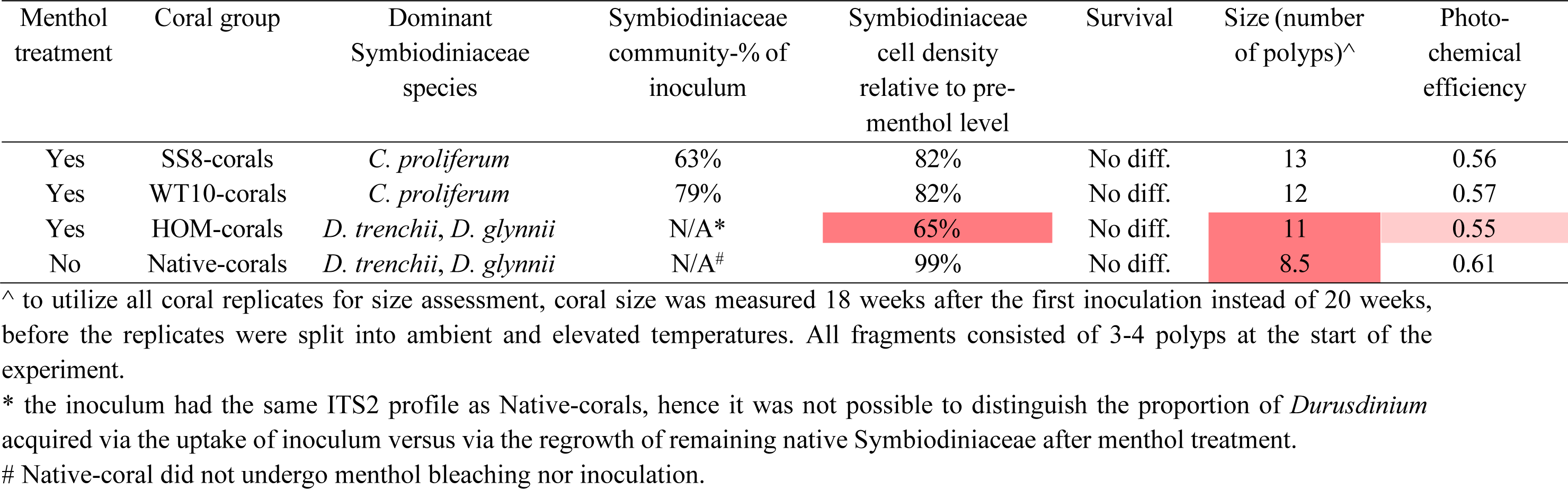
Summary of inoculation outcome and physiological performance of coral groups under ambient temperature, 20 weeks after the first inoculation. Dark red indicates a strong/moderate negative effect, whereas light red indicates a mild (≤ 10% change) negative effect.

**Table 4.** Comparison of the Symbiodiniaceae community, physiological performance and metabolite profile within coral groups under ambient versus elevated temperatures, at the end of the short-term heat stress experiment (i.e., 8 days exposure to 32°C). Dark red indicates a strong/moderate negative effect, whereas light red indicates a mild (≤ 10% change) negative effect.

| Menthol treatment | Coral group | Dominant Symbiodiniaceae species | Symbiodiniaceae community change | Symbiodiniaceae cell density difference | Survival difference | Photo-chemical efficiency difference | No. of metabolites of significantly different intensities |
| --- | --- | --- | --- | --- | --- | --- | --- |
| Yes | SS8-corals | *C. prolifera* | No change | 0% | +3% | No change | 12 |
| Yes | WT10-corals | *C. prolifera* | -21% of WT10 | -89% | -12% | -44% | 66 |
| Yes | HOM-corals | *D. trenchii*, *D. glynnii* | No change | 0% | No change | No change | 0 |
| No | Native-corals | *D. trenchii*, *D. glynnii* | No change | -10% | No change | -5% | 0 |

### 3.4 Long-term monitoring and mock community

Heat-evolved *C. proliferum* (SS8) was detected in all SS8-corals with moderate relative abundance one year (averaged 32%) and two years (averaged 42%) after its first inoculation, suggesting long-stability of this novel symbiosis (Fig. 11).

**Fig. 11.**
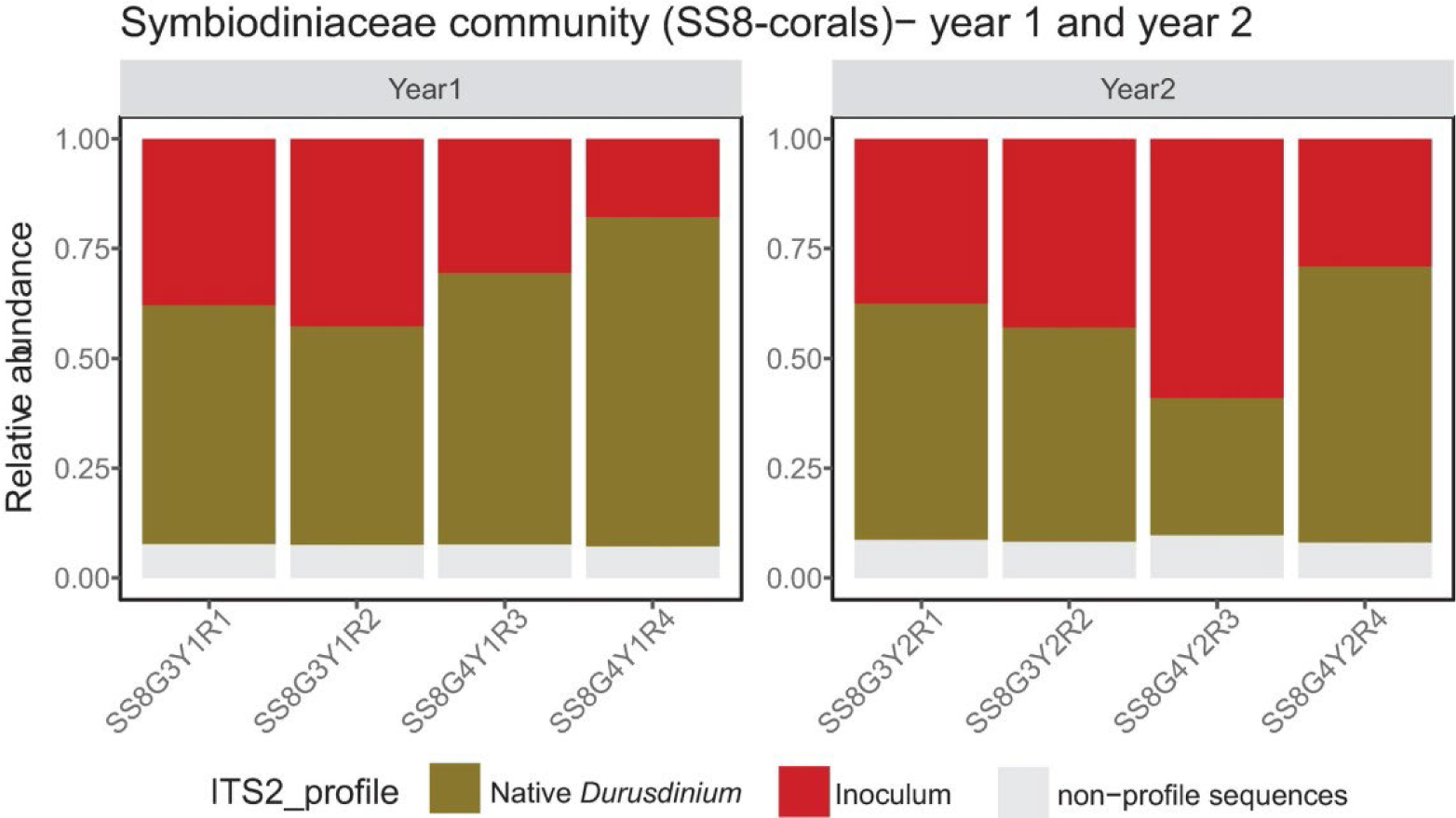
Coral-associated Symbiodiniaceae communities of SS8-corals one year and two years after the first inoculation of heat-evolved *C. proliferum* (SS8 strain). “G” refers to the sample’s genotype, “R refers to the sample’s replicate number.

## 4. DISCUSSION

### 4.1 Heat-evolved *C. proliferum* can form a symbiosis with adult corals despite being a heterologous symbiont

While heat-evolved *C. proliferum* (SS8) has been experimentally evolved (Chakravarti et al. 2017) and maintained at 31°C for ∼10 years, it can form a symbiosis with adult GBR *G. fascicularis*, to which it is a heterologous symbiont. SS8 can also colonize aposymbiotic coral larvae of a taxonomically distant species (*Acropora tenuis*) (Buerger et al. 2020), indicating it can potentially be a reef restoration resource applicable to corals across taxa and life stages. Consistent with previous studies (Wang et al. 2012; Scharfenstein et al. 2022; Puntin et al. 2023), menthol treatment was an effective way to produce corals with minimal native Symbiodiniaceae (< 1-2% of pre-bleaching densities), but cannot produce fully aposymbiotic corals. Nevertheless, the presence of remnant native *Durusdinium* sp. did not prevent the uptake of heterologous *C. proliferum* (both wild-type and heat-evolved strains) in adult *G. fascicularis*. Although the inocula only accounted for a small proportion of the endosymbiotic Symbiodiniaceae communities seven weeks after the first inoculation, they continued to populate the corals, achieving a moderate to high relative abundance (43-82%) by 20 weeks. This suggests that the generalist species, *C. proliferum*, is compatible with *G. fascicularis*. While SS8 and WT10 originated from the same mother culture from 10 years ago, SS8 was taken up by two of the four coral genotypes tested but WT10 was only taken up by one genotype. The result indicates intraspecific host genotypic effects play a role in symbiont uptake.

The inoculation outcome of WT10 was consistent with that of Scharfenstein et al. (2022) on six other adult coral species, all of which formed a symbiosis with WT10 at consistently high relative abundance. However, note that the above study only used one coral genotype per coral species, therefore host genotypic effects could not be evaluated. The proportion of WT10 in *G. fascicularis* was fairly homogeneous among replicates, but the relative abundance of SS8 varied across replicates from the same coral genotype under both temperatures. This contrasting pattern may have been a consequence of different growth rates between WT10 and SS8, which could have determined the competition outcome with native *Durusdinium* sp.. Under ambient temperature, *in vitro* growth rates are higher for *C. proliferum* than *D. trenchii* or *D. glynnii* (C. Alvarez-Roa, personal communication, 16^th^ Jun 2023), and WT10 grows faster than SS8 (Buerger et al. 2020). Assuming growth rate differences persist *in hospite*, WT10 was likely able to colonize the corals with little competition from the remnant, more slowly growing native *Durusdinium* sp., resulting in a consistently high relative abundance among replicates. SS8, in contrast, had less of an advantage in growth rate and was therefore likely experiencing stronger competition with native *Durusdinium* sp., resulting in a less predictable and less consistent inoculation outcome. No such contrast in replicate variability was found in aposymbiotic coral larvae inoculated with WT10 or SS8, where competition from native symbionts was absent (Buerger et al. 2020).

Despite the inoculation success observed here, a failure of symbiosis establishment between bleached adult corals and exogenously supplied, laboratory cultured, heterologous Symbiodiniaceae has been reported in several earlier studies (Coffroth et al. 2010; Morgans et al. 2020). Following a thermal treatment that reduced > 98% of the corals’ native Symbiodiniaceae population and the inoculation with heterologous strains of *Symbiodinium*, *Breviolum* and *Durusdinium*, only *Breviolum* was detected in some adult *Porites divaricata* three weeks post inoculation (Coffroth et al. 2010). This inoculum was no longer detected in *P. divaricata* by five weeks post inoculation. One possible explanation of this contrast is the low inoculation cell density and the absence of feed during inoculation. This study and Scharfenstein et al. (2022) employed ∼10^4^ cell mL^-1^ (i.e., 10 times more than in Coffroth et al. (2010)) and added food (*Artemia nauplii* or microalgae) during inoculation to encourage phagocytosis. Given that these adult corals were not aposymbiotic and contained remnant native Symbiodiniaceae, a high inoculation cell density and food supply could be vital for successful colonization by exogenous Symbiodiniaceae. In another study, thermally bleached *Acropora millepora* (naturally dominated by Cspc, C3k) was inoculated with heterologous *D. trenchii* and *C. proliferum* culture and the inoculation was also unsuccessful (Morgans et al. 2020). However, their inoculation was conducted under elevated temperature (32°C), and it has previously been shown that symbiont uptake is limited under elevated temperatures (Abrego et al. 2012). Given the significant potential of adult coral inoculation experiments in designing tools for reef restoration, testing parameters to achieve optimal inoculation success will be invaluable in future studies.

### 4.2 *Cladocopium proliferum* accelerated recovery post menthol treatment

While SS8-, WT10- and HOM-corals underwent the same menthol treatment and inoculation process, the two groups dominated by *C. proliferum* (SS8- and WT10-corals) recovered faster than the group dominated by *D. trenchii* and *D. glynnii* (HOM-corals) (note that recovery occurred under ambient temperature). This is evidenced by higher Symbiodiniaceae cell density (SS8- and WT10-corals), coral size (SS8-corals) and photochemical efficiency (WT10-corals) compared to HOM-corals (Table 3). While these corals were inoculated with the same Symbiodiniaceae cell density, SS8- and WT10-corals had recovered to ∼82% of pre-menthol treatment level cell density, while HOM-coral only recovered to ∼65% by 20 weeks after the end of menthol treatment. Several studies have also shown the advantages of symbiosis with *Cladocopium* under favorable, ambient temperatures on the GBR. For example, *A. tenuis* juveniles dominated by *Cladocopium* C1 grew 2-3 times faster in the field than those dominated by *Durusdinium* D1 (Little et al. 2004). Adult *A. millepora* dominated by *Durusdinium* sp. also grew 29 and 38% slower than conspecifics dominated by *Cladocopium* C2 in the lab and field respectively (Jones and Berkelmans 2010). Based on these findings, the inoculation with laboratory cultured *C. proliferum* is a possible tool to accelerate coral recovery following a mass bleaching event, as long as the seawater temperature has returned to and continues to stay at favorable ambient condition. However, how will these *Cladocopium*-dominated corals perform if the reef experiences yet another marine heatwave?

### 4.3 Heat-evolved Symbiodiniaceae conferred high bleaching tolerance to adult corals

Our findings showed corals hosting the wild-type *C. proliferum* were heat-sensitive while corals hosting the heat-evolved strain SS8 were as tolerant to elevated temperature as HOM-corals. These results demonstrate that heat-evolved *C. proliferum* can confer a similar level of thermal tolerance to *G. fascicularis* as the naturally thermally tolerant *D. trenchii* and *D. glynnii*. In contrast, WT10- corals underwent 12% mortality, a 44% drop in photochemical efficiency and a 89% decline in Symbiodiniaceae cell density (Table 4). Under elevated temperature, WT10-corals shifted toward a more *Durusdinium*-dominated community and reduced the proportion of WT10 by 21%. *G. fascicularis* may have preferentially removed WT10 due to its high level of reactive oxygen species released (ROS) (Buerger et al. 2020) and its lower carbon acquisition ability than *Durusdinium* sp. under elevated temperature (Baker et al. 2013). Alternatively, WT10 may have had a higher mortality rate under elevated temperature.

Shuffling and/or switching to *Durusdinium* sp. is a commonly reported response to elevated temperatures and is interpreted as an adaptive response to ocean warming (Chen et al. 2005; Berkelmans and van Oppen 2006; Mieog et al. 2007; Stat et al. 2013; Silverstein et al. 2015; Boulotte et al. 2016). Heat-evolved *C. proliferum* may be superior to wild-type *D. trenchii* and *D. glynnii* as it recovers to pre-bleaching densities faster when the temperature is benign and confers higher coral growth rates. The metabolite data confirmed that some of the SS8-coral replicates were metabolically more similar to *Durusdinium*-dominated coral under elevated temperature. The physiological data also aligns closely with the corals’ metabolite profiles, with WT10-corals showing the most significant difference between ambient and elevated temperatures. No metabolite profile change was observed in *Durusdinium*-dominated corals under elevated temperature, as expected, and this was consistent with the physiological data where no (HOM-corals) or limited (Native-corals) negative effects were observed. The enhanced thermal tolerance of SS8-corals did not show a trade-off with coral growth under ambient temperature, which is typically found in *Durusdinium*-dominated corals from the GBR (Little et al. 2004; Jones and Berkelmans 2010; Cunning et al. 2015). A considerable level of natural variation in heat tolerance within a population of *Acropora digitifera* which hosts exclusively *Cladocopium* C40 also occurred without a trade-off against growth; in fact, a positive correlation between heat tolerance and growth was observed (Lachs et al. 2023). This suggests that the selection for heat tolerance does not necessarily compromise coral growth.

### 4.4 Implications of contrasting metabolite profiles between heat-evolved and wild-type holobionts

Nearly 40% of the metabolites that significantly differed in relative intensities between WT10- and SS8-corals under elevated temperature are pigments, co-localized with the algal symbionts. These are minor carotenoid-like pigments and can be found co-varying with chlorophyll levels. Carotenoid pigments are generally biosynthesized by autotrophic marine organisms (e.g., algae, and some bacteria, archaea and fungi). They are known to protect the photosynthetic machinery in a number of ways, including 1) limiting lipid peroxidation, 2) scavenging singlet oxygen (^1^O_2_), 3) preventing the formation of ^1^O_2_ by reacting with chlorophyll in its triplet state (^3^Chl), and 4) dissipating the excess excitation energy formed through the xanthophyll cycle (Das and Roychoudhury 2014; Galasso et al. 2017). Carotenoid-like pigments mostly protect lipid membranes (by their associated lipophilicity). The contrasting algal pigment profiles of WT10- and SS8-corals suggests that a difference in photoprotective mechanisms may contribute to the enhanced thermal tolerance observed in SS8-corals.

Another notable change in the metabolite pool is a downregulation of seven ceramide species in WT10-corals. Ceramides are intermediates for the biosynthesis and metabolism of sphingolipids. This lipid class is involved in cell signaling, regulates the balance between apoptosis, cell survival and proliferation (Rosset et al. 2021; Bhattacharya 2022); and several lines of evidence have pointed towards its regulatory role in the cnidarian-Symbiodiniaceae symbiosis (Rodriguez-Lanetty et al. 2006; Detournay and Weis 2011; Kitchen and Weis 2017; Kitchen et al. 2017). Its downregulation in WT10-coral suggested that acute heat stress may have caused the corals to stop acquiring exogenous Symbiodiniaceae and to increase algal symbiont expulsion to avoid further cellular damage by ROS.

Both *Cladocopium*-dominated groups (WT10- and SS8-corals) showed downregulation of PC, Cer and betaine lipids under elevated temperature, suggesting that these metabolite classes could be a *Cladocopium*-specific response to high temperatures. The localization of PC and betaine lipids are consistent with our previous study (Chan et al. 2023). Nevertheless, SS8-corals experienced much smaller fold changes and a much lower number of significantly different metabolites. Compared to SS8, a higher level of secreted reactive oxygen species (ROS) has been detected in WT10 *in vitro* under elevated temperature (Buerger et al. 2020). WT10-corals therefore likely experienced higher levels of ROS than SS8-croals under elevated temperature, although this was not measured in this study. As ROS indiscriminately reacts with some metabolites, proteins and polyunsaturated fatty acyl chains via peroxidation, the elevated temperature experienced by WT10-corals could result in a larger and mixed altered pool of metabolites. To test this hypothesis, future studies will have to conduct a targeted analysis on the products of ROS oxidation, such as reactive carbonyl species (Yalcinkaya et al. 2019).

### 4.5 Equilibrium symbiont levels were not reached over a 20-week period

Despite a long (20 weeks) recovery period following symbiont inoculation, none of the menthol-treated coral groups had recovered pre-treatment Symbiodiniaceae cell densities and had slightly lower photochemical efficiency than control (non-menthol-treated) corals. HOM-corals were inoculation with homologous Symbiodiniaceae and should achieve the same cell density and photochemical efficiency as Native-corals when fully recovered, whereas SS8- and WT10-corals may arrive to a different equilibrium level when fully recovered given they are a different species to the native symbionts. Nevertheless, it is worth noting that the dark-adapted yields of the menthol-treated coral groups (0.55-0.57) under ambient temperature were within the healthy range for corals.

While menthol is widely used to remove native Symbiodiniaceae in the coral model sea anemone, *Exaiptasia diaphana* (‘Aiptasia’) (e.g., Matthews et al. 2016, 2017; Gabay et al. 2019; Tortorelli et al. 2020; Tsang Min Ching et al. 2022), and more recently in adult corals (e.g., Scharfenstein et al. 2022; Puntin et al. 2023), little is known of its potential long-term effect on cnidarians. Menthol is a cyclic terpene alcohol and has been hypothesized to cause coral bleaching via either photoinhibition of Symbiodiniaceae (Wang et al. 2017) or via Ca^2+^-triggered exocytosis by binding to the transient receptor potential TRPM8 (Pang and Südhof 2010). Matthews et al. (2016) reported no difference in survival, photosynthesis and respiration performance between menthol-treated and inoculated Aiptasia and untreated symbiotic anemones by 12 weeks. In the coral, *Isopora palifera*, a significant reduction (44-50%) in glutamate dehydrogenase activity, total free amino acids and essential free amino acids, as well as a change in dominant free amino acid species, were found in adults 6 to 10 days after menthol treatment, reflecting major effects on the holobiont’s nitrogen metabolism (Wang et al. 2012). In contrast, no such change was observed in *Stylophora pistillata* that underwent the same menthol treatment. However, these observations were based on freshly bleached corals and no longer-term monitoring has been conducted.

Our data detected the legacy of menthol treatment 20 weeks after its cessation. MSI results show a clear separation between the menthol-treated and control groups under ambient temperature. Among the top significant metabolites, ceramide species stood out as one of the key contributing groups. Compared to control corals, all menthol-treated/inoculated groups had consistently higher relative intensity in these ceramide species, with some HOM-coral replicates showing particular high intensity. Higher ceramide relative intensity, together with the fact that these corals have not recovered to pre-treatment level cell density, suggests that these corals could still be in the active process of regulating and incorporating exogenous Symbiodiniaceae. In support of this notion, the highest ceramide relative intensity was found in HOM-corals which had the lowest cell density.

### 4.6 Implications for reef restoration, limitations and future directions

We have demonstrated for the first time that heat-evolved Symbiodiniaceae can form a symbiosis with adult corals and that this novel symbiotic relation continues for two years. Heat-evolved *C. proliferum* accelerated coral recovery from chemical bleaching under ambient temperature, and conferred thermal bleaching tolerance to adult corals to a similar extent of that of the naturally heat tolerant *D. trenchii* and *D. glynnii*. Since heat-evolved *C. proliferum* can also form a symbiosis with and confer thermotolerance to coral larvae and recruits of a taxonomically distant species (Buerger et al. 2020), these symbionts are a potentially valuable resource for reef restoration applicable across coral species and life stages. Enhanced thermotolerance will ‘buy time’ for reefs until carbon emission is curtailed and the climate is stabilized. The long-term stability of the symbiosis with heat-evolved *C. proliferum* offers hope that this symbiont population could self-sustain and may continue to provide benefits to the coral hosts for many years. However, it is also possible that heat-evolved *C. proliferum* could lose its thermotolerance after long-term exposure to ambient temperature, and this is an important avenue for future studies. We acknowledge that this study was laboratory-based using only one coral species and one experimentally evolved Symbiodiniaceae strain and recommend future studies to: 1) expand to multiple coral species, 2) utilize multiple experimentally evolved Symbiodiniaceae strains, 3) test holobiont performance in the field. To maximize the success of utilizing heat-evolved Symbiodiniaceae in reef restoration, the development of best practice protocols (e.g., optimal inoculum cell density, temperature, type of food supplement and extent of menthol bleaching) are essential.

## ACKNOWLEDGEMENT

We thank J. Ahern for aquarium support; the National Sea Simulator team, especially A. Severati and L. Koukoumaftsis for coral collection; M. Nitschke for the Symbiodiniaceae community statistical pipeline and S. J. Tsang Min Ching for assistance in coral inoculation and library preparation. Thanks are also extended to D. Baker, S. McIlroy and G. Puntin for fruitful discussions and protocol sharing on coral menthol bleaching; as well as the Metabolomics Australia, especially V. Lui and V. Narayana for technical support. This research was funded by the Australian Research Council Laureate Fellowship to MJHvO (FL180100036), the Paul G. Allen Family Foundation, and the Reef Restoration and Adaptation Program- a partnership between the Australian Governments Reef Trust and the Great Barrier Reef Foundation.

## DATA AVAILABILITY

All raw data and R codes for statistical analysis are available on https://doi.org/10.5281/zenodo.8008851. Details of each file are provided in Table S15. Metabolite annotations on MetaSpace are available at https://metaspace2020.eu/ (project name: heat-evolved algal symbionts enhance thermotolerance of adult corals without growth trade-off). Raw sequences of the ITS2 Symbiodiniaceae data sets are available in GenBank (project Accession no.: PRJNA983409) (embargoed access).

## AUTHOR CONTRIBUTIONS

WY Chan: conceptualization, data curation, formal analysis, validation, investigation, visualization, methodology, supervision, project administration, and writing—original draft, review, and editing. D Rudd: resources, data curation, software, formal analysis, validation, investigation, visualization, methodology, and writing—review and editing. L Meyers: data curation, formal analysis, investigation, visualization, and writing—review and editing. SH Topa: data curation and writing—review and editing. MJH van Oppen: conceptualization, resources, supervision, funding acquisition, project administration, and writing—original draft, review, and editing.

## REFERENCES

Abrego D, Willis BL, van Oppen MJH (2012) Impact of light and temperature on the uptake of algal symbionts by coral juveniles. PLOS ONE 7:e50311

Allen-Waller L, Barott KL (2023) Symbiotic dinoflagellates divert energy away from mutualism during coral bleaching recovery. Symbiosis 89:173–186

Anderson MJ, Ellingsen KE, McArdle BH (2006) Multivariate dispersion as a measure of beta diversity. Ecology Letters 9:683–693

Baker AC, Starger CJ, McClanahan TR, Glynn PW (2004) Corals’ adaptive response to climate change. Nature 430:741–741

Baker DM, Andras JP, Jordán-Garza AG, Fogel ML (2013) Nitrate competition in a coral symbiosis varies with temperature among *Symbiodinium* clades. ISME J 7:1248–1251

Benjamini Y, Hochberg Y (1995) Controlling the false discovery rate: a practical and powerful approach to multiple testing. Journal of the Royal Statistical Society Series B (Methodological) 57:289–300

Berkelmans R, van Oppen MJH (2006) The role of zooxanthellae in the thermal tolerance of corals: a ‘nugget of hope’ for coral reefs in an era of climate change. Proceedings of the Royal Society of London B: Biological Sciences 273:2305–2312

Bhattacharya A (2022) Lipid metabolism in plants under low-temperature stress: a review. In: Bhattacharya A. (eds) Physiological processes in plants under low temperature stress. Springer, Singapore, pp 409–516

Bindoff N, Cheung W, Kairo J, Arístegui J, Guinder V, Hallberg R, Hilmi N, Jiao N, Karim M, Levin L, O’Donoghue S, Cuicapusa Purca S, Rinkevich B, Suga T, Tagliabue A, Williamson P (2019) Changing ocean, marine ecosystems, and dependent communities. IPCC special report on the ocean and cryosphere in a changing climate. [H-O Pörtner, DC Roberts, V Masson-Delmotte, P Zhai, M Tignor, E Poloczanska, K Mintenbeck, A Alegría, M Nicolai, A Okem, J Petzold, B Rama, NM Weyer (eds.)]. Cambridge: University Press, pp 477–587

Bolker BM, Brooks ME, Clark CJ, Geange SW, Poulsen JR, Stevens MHH, White J-SS (2009) Generalized linear mixed models: a practical guide for ecology and evolution. Trends in Ecology & Evolution 24:127–135

Boulotte NM, Dalton SJ, Carroll AG, Harrison PL, Putnam HM, Peplow LM, van Oppen MJH (2016) Exploring the *Symbiodinium* rare biosphere provides evidence for symbiont switching in reef-building corals. ISME J 10:2693–2701

Buerger P, Alvarez-Roa C, Coppin CW, Pearce SL, Chakravarti LJ, Oakeshott JG, Edwards OR, Oppen MJH van (2020) Heat-evolved microalgal symbionts increase coral bleaching tolerance. Science Advances 6:eaba2498

Butler CC, Turnham KE, Lewis AM, Nitschke MR, Warner ME, Kemp DW, Hoegh-Guldberg O, Fitt WK, van Oppen MJH, LaJeunesse TC (2023) Formal recognition of host-generalist species of dinoflagellate (*Cladocopium*, Symbiodiniaceae) mutualistic with Indo-Pacific reef corals. Journal of Phycology n/a:

Camp EF, Edmondson J, Doheny A, Rumney J, Grima AJ, Huete A, Suggett DJ (2019) Mangrove lagoons of the Great Barrier Reef support coral populations persisting under extreme environmental conditions. Marine Ecology Progress Series 625:1–14

Cantin NE, van Oppen MJH, Willis BL, Mieog JC, Negri AP (2009) Juvenile corals can acquire more carbon from high-performance algal symbionts. Coral Reefs 28:405

Chakravarti LJ, Beltran VH, van Oppen MJH (2017) Rapid thermal adaptation in photosymbionts of reef-building corals. Global Change Biology 23:4675–4688

Chan WY, Oakeshott JG, Buerger P, Edwards OR, Oppen MJH van (2021) Adaptive responses of free-living and symbiotic microalgae to simulated future ocean conditions. Global Change Biology 27:1737–1754

Chan WY, Rudd D, Oppen MJ van (2023) Spatial metabolomics for symbiotic marine invertebrates. Life Science Alliance 6:

Chen CA, Wang J-T, Fang L-S, Yang Y-W (2005) Fluctuating algal symbiont communities in *Acropora palifera* (Scleractinia: Acroporidae) from Taiwan. Marine Ecology Progress Series 295:113–121

Coffroth MA, Poland DM, Petrou EL, Brazeau DA, Holmberg JC (2010) Environmental symbiont acquisition may not be the solution to warming seas for reef-building corals. PLOS ONE 5:e13258

Cribari-Neto F, Zeileis A (2010) Beta regression in R. Journal of Statistical Software 34:1–24

Cunning R, Gillette P, Capo T, Galvez K, Baker AC (2015) Growth tradeoffs associated with thermotolerant symbionts in the coral *Pocillopora damicornis* are lost in warmer oceans. Coral Reefs 34:155–160

Das K, Roychoudhury A (2014) Reactive oxygen species (ROS) and response of antioxidants as ROS-scavengers during environmental stress in plants. Frontiers in Environmental Science 2:

Davy SK, Allemand D, Weis VM (2012) Cell biology of cnidarian-dinoflagellate symbiosis. Microbiology and Molecular Biology Reviews 76:229–261

Detournay O, Weis VM (2011) Role of the sphingosine rheostat in the regulation of cnidarian-dinoflagellate symbioses. The Biological Bulletin 221:261–269

Dunn OJ (1964) Multiple comparisons using rank sums. Technometrics 6:241–252

Gabay Y, Parkinson JE, Wilkinson SP, Weis VM, Davy SK (2019) Inter-partner specificity limits the acquisition of thermotolerant symbionts in a model cnidarian-dinoflagellate symbiosis. ISME J 13:2489–2499

Galasso C, Corinaldesi C, Sansone C (2017) Carotenoids from marine organisms: biological functions and industrial applications. Antioxidants (Basel) 6:96

Hoegh-Guldberg O (1999) Climate change, coral bleaching and the future of the world’s coral reefs. Mar Freshwater Res 50:839–866

Hughes TP, Anderson KD, Connolly SR, Heron SF, Kerry JT, Lough JM, Baird AH, Baum JK, Berumen ML, Bridge TC, Claar DC, Eakin CM, Gilmour JP, Graham NAJ, Harrison H, Hobbs J-PA, Hoey AS, Hoogenboom M, Lowe RJ, McCulloch MT, Pandolfi JM, Pratchett M, Schoepf V, Torda G, Wilson SK (2018) Spatial and temporal patterns of mass bleaching of corals in the Anthropocene. Science 359:80

Hume BCC, Smith EG, Ziegler M, Warrington HJM, Burt JA, LaJeunesse TC, Wiedenmann J, Voolstra CR (2019) SymPortal: A novel analytical framework and platform for coral algal symbiont next-generation sequencing ITS2 profiling. Molecular Ecology Resources 19:1063–1080

Jones A, Berkelmans R (2010) Potential costs of acclimatization to a warmer climate: growth of a reef coral with heat tolerant vs. sensitive symbiont types. PLOS ONE 5:e10437

Jones AM, Berkelmans R (2011) Tradeoffs to thermal acclimation: energetics and reproduction of a reef coral with heat tolerant *Symbiodinium* type-D. Journal of Marine Biology 2011:e185890

Kitchen SA, Poole AZ, Weis VM (2017) Sphingolipid metabolism of a sea anemone is altered by the presence of dinoflagellate symbionts. The Biological Bulletin 233:242–254

Kitchen SA, Weis VM (2017) The sphingosine rheostat is involved in the cnidarian heat stress response but not necessarily in bleaching. Journal of Experimental Biology 220:1709–1720

Lachs L, Humanes A, Pygas DR, Bythell JC, Mumby PJ, Ferrari R, Figueira WF, Beauchamp E, East HK, Edwards AJ, Golbuu Y, Martinez HM, Sommer B, van der Steeg E, Guest JR (2023) No apparent trade-offs associated with heat tolerance in a reef-building coral. Commun Biol 6:1–10

LaJeunesse TC (2005) “Species” radiations of symbiotic dinoflagellates in the Atlantic and Indo-Pacific since the miocene-pliocene transition. Molecular Biology and Evolution 22:570– 581

Little AF, van Oppen MJH, Willis BL (2004) Flexibility in algal endosymbioses shapes growth in reef corals. Science 304:1492

Matthews JL, Crowder CM, Oakley CA, Lutz A, Roessner U, Meyer E, Grossman AR, Weis VM, Davy SK (2017) Optimal nutrient exchange and immune responses operate in partner specificity in the cnidarian-dinoflagellate symbiosis. PNAS 114:13194–13199

Matthews JL, Sproles AE, Oakley CA, Grossman AR, Weis VM, Davy SK (2016) Menthol-induced bleaching rapidly and effectively provides experimental aposymbiotic sea anemones (*Aiptasia* sp.) for symbiosis investigations. J Exp Biol 219:306

Mieog JC, van Oppen MJH, Cantin NE, Stam WT, Olsen JL (2007) Real-time PCR reveals a high incidence of *Symbiodinium* clade D at low levels in four scleractinian corals across the Great Barrier Reef: implications for symbiont shuffling. Coral Reefs 26:449–457

Morgans CA, Hung JY, Bourne DG, Quigley KM (2020) Symbiodiniaceae probiotics for use in bleaching recovery. Restoration Ecology 28:282–288

Muller-Parker G, D’Elia CF, Cook CB (2015) Interactions between corals and their symbiotic algae. In: Birkeland C. (eds) Coral Reefs in the Anthropocene. Springer Netherlands, Dordrecht, pp 99–116

Oksanen J, Blanchet FG, Friendly M, Kindt R, Legendre P, McGlinn D, Minchin PR, O’Hara RB (2016) Vegan: community ecology package. R package version 2.4–1.

Oliver TA, Palumbi SR (2011) Do fluctuating temperature environments elevate coral thermal tolerance? Coral Reefs 30:429–440

Ortiz JC, González-Rivero M, Mumby PJ (2013) Can a thermally tolerant symbiont improve the future of Caribbean coral reefs? Global Change Biology 19:273–281

Palacio-Castro AM, Smith TB, Brandtneris V, Snyder GA, van Hooidonk R, Maté JL, Manzello D, Glynn PW, Fong P, Baker AC (2023) Increased dominance of heat-tolerant symbionts creates resilient coral reefs in near-term ocean warming. Proceedings of the National Academy of Sciences 120:e2202388120

Pang ZP, Südhof TC (2010) Cell biology of Ca^2+^-triggered exocytosis. Current Opinion in Cell Biology 22:496–505

Puntin G, Craggs J, Hayden R, Engelhardt KE, McIlroy S, Sweet M, Baker DM, Ziegler M (2023) The reef-building coral *Galaxea fascicularis*: a new model system for coral symbiosis research. Coral Reefs 42:239–252

Quigley KM, Baker AC, Coffroth MA, Willis BL, van Oppen MJH (2018) Bleaching resistance and the role of algal endosymbionts. In: van Oppen M.J.H., Lough J.M. (eds) Coral bleaching: patterns, processes, causes and consequences. Springer International Publishing, Cham, pp 111–151

Quigley KM, van Oppen MJH (2022) Predictive models for the selection of thermally tolerant corals based on offspring survival. Nat Commun 13:1543

Quigley KM, Ramsby B, Laffy P, Harris J, Mocellin VJL, Bay LK (2022) Symbioses are restructured by repeated mass coral bleaching. Science Advances 8:eabq8349

Roach TNF, Dilworth J, H CM, Jones AD, Quinn RA, Drury C (2021) Metabolomic signatures of coral bleaching history. Nature Ecology & Evolution 1–9

Rodriguez-Lanetty M, Phillips WS, Weis VM (2006) Transcriptome analysis of a cnidarian – dinoflagellate mutualism reveals complex modulation of host gene expression. BMC Genomics 7:23

Rosset SL, Oakley CA, Ferrier-Pagès C, Suggett DJ, Weis VM, Davy SK (2021) The molecular language of the cnidarian–dinoflagellate symbiosis. Trends in Microbiology 29:320–333

Scharfenstein HJ, Chan WY, Buerger P, Humphrey C, van Oppen MJH (2022) Evidence for de novo acquisition of microalgal symbionts by bleached adult corals. The ISME Journal

Silverstein RN, Cunning R, Baker AC (2015) Change in algal symbiont communities after bleaching, not prior heat exposure, increases heat tolerance of reef corals. Global Change Biology 21:236–249

Stat M, Gates RD (2011) Clade D *Symbiodinium* in scleractinian corals: a “nugget” of hope, a selfish opportunist, an ominous sign, or all of the above? Journal of Marine Biology 2011:1–9

Stat M, Pochon X, Franklin EC, Bruno JF, Casey KS, Selig ER, Gates RD (2013) The distribution of the thermally tolerant symbiont lineage (*Symbiodinium* clade D) in corals from Hawaii: correlations with host and the history of ocean thermal stress. Ecology and Evolution 3:1317–1329

Tortorelli G, Belderok R, Davy SK, McFadden GI, van Oppen MJH (2020) Host genotypic effect on algal symbiosis establishment in the coral model, the anemone *Exaiptasia diaphana*, from the Great Barrier Reef. Frontiers in Marine Science 6:833

Tsang Min Ching SJ, Chan WY, Perez-Gonzalez A, Hillyer KE, Buerger P, van Oppen MJH (2022) Colonization and metabolite profiles of homologous, heterologous and experimentally evolved algal symbionts in the sea anemone *Exaiptasia diaphana*. ISME COMMUN 2:1–10

Wang J-T, Chen Y-Y, Tew KS, Meng P-J, Chen CA (2012) Physiological and biochemical performances of menthol-induced aposymbiotic corals. PLOS ONE 7:e46406

Wang J-T, Keshavmurthy S, Chu T-Y, Chen CA (2017) Diverse responses of Symbiodinium types to menthol and DCMU treatment. PeerJ 5:e3843

Wepfer PH, Nakajima Y, Hui FKC, Mitarai S, Economo EP (2020) Metacommunity ecology of Symbiodiniaceae hosted by the coral *Galaxea fascicularis*. Marine Ecology Progress Series 633:71–87

Yalcinkaya T, Uzilday B, Ozgur R, Turkan I, Mano J (2019) Lipid peroxidation-derived reactive carbonyl species (RCS): Their interaction with ROS and cellular redox during environmental stresses. Environmental and Experimental Botany 165:139–149

