## Supplementary Information for "Heat-evolved algal symbionts enhance bleaching tolerance of adult corals without trade-off against growth"

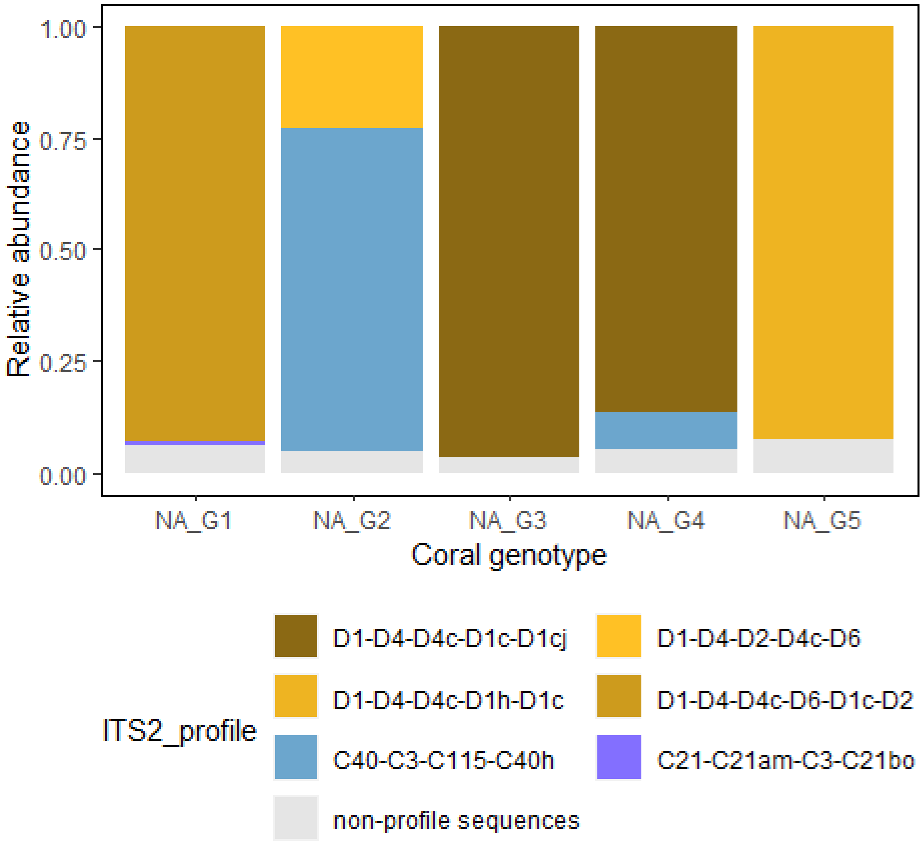

Fig. S1. The native coral-associated Symbiodiniaceae communities of the five *G. fascicularis* colonies (G1-G5) of this study.

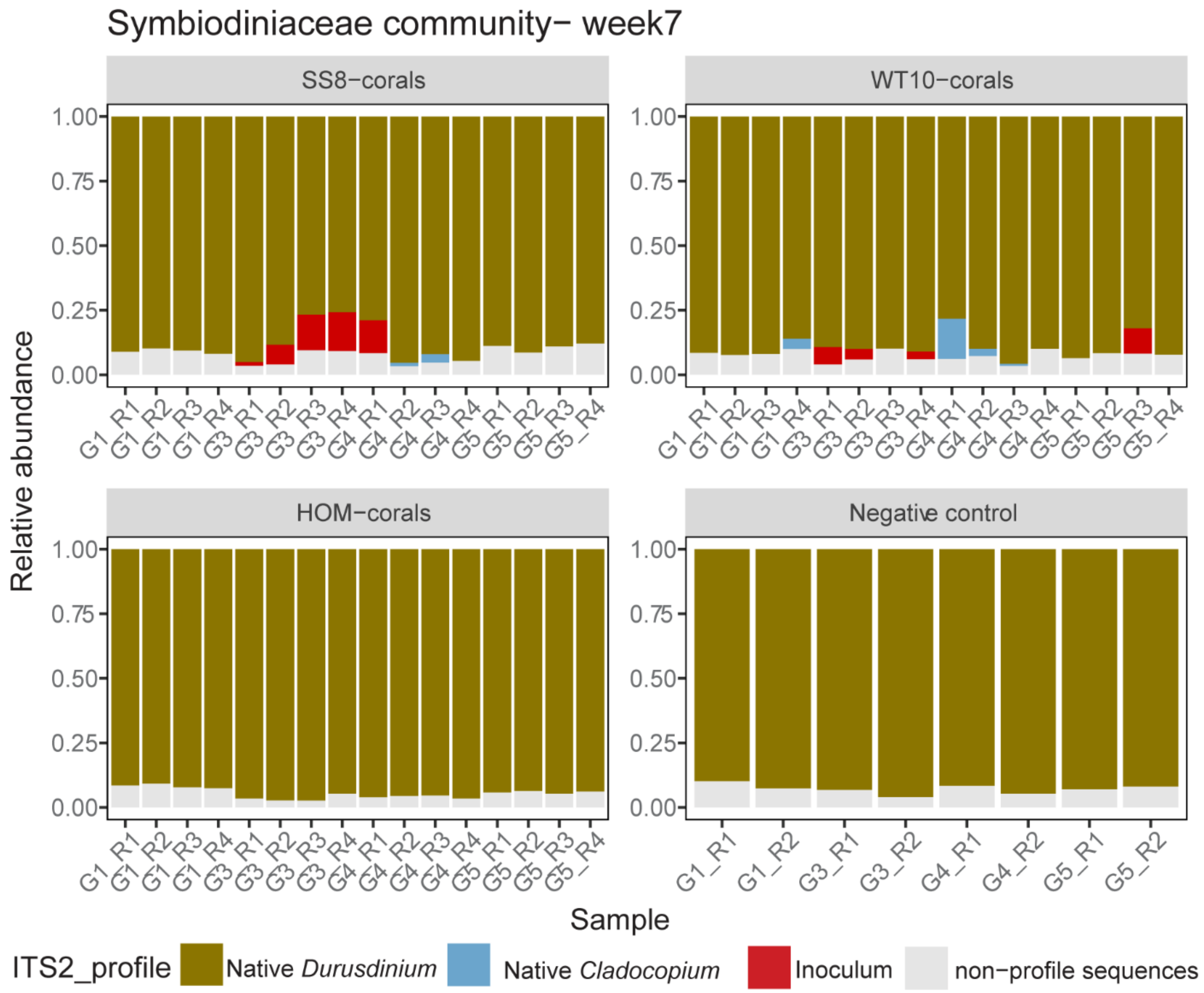

Fig. S2. Coral-associated Symbiodiniaceae communities seven weeks after the first inoculation. Note that the corals were inoculated three times (at week 0, week 4, week 7) and these samples were taken before the week 7 inoculation. The abbreviations of the experimental groups refer to: corals that were inoculated and dominated by heat-evolved *Cladocopium proliferum* (SS8-corals), wild-type *C. proliferum* (WT10-corals), freshly isolated, homologous Symbiodiniaceae from *G. fascicularis* (HOM-corals); as well as control corals with no menthol treatment nor inoculation (Native-corals). “G” refers to the sample’s genotype, “R refers to the sample’s replicate number.

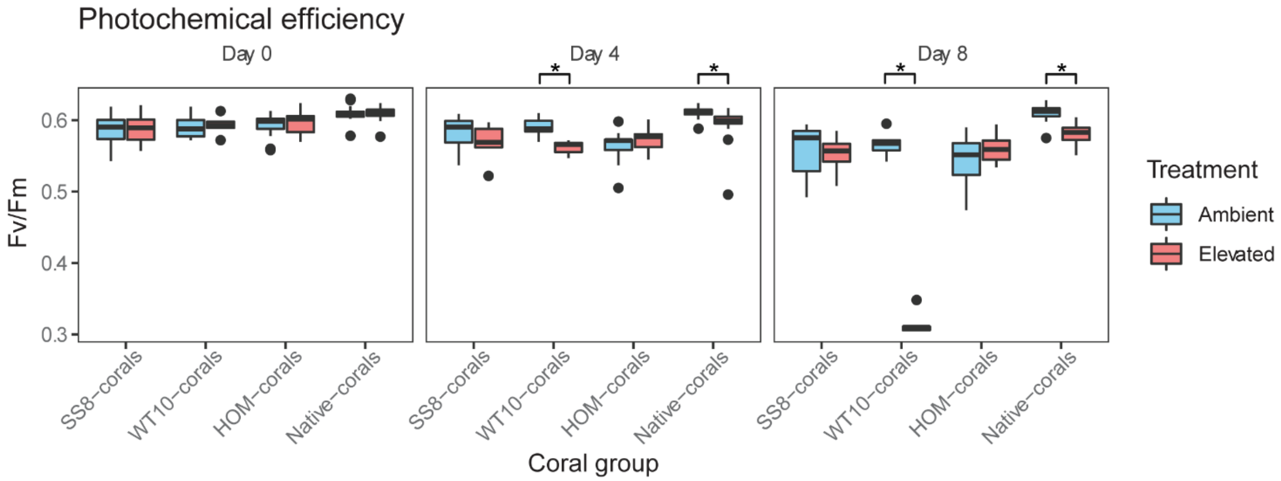

Fig. S3. Coral photochemical efficiency (maximum quantum yield, Fv/Fm) before temperature ramping (day 0), on the 4$^{th}$ day since exposure to 32°C (day 4) and on the 8$^{th}$ day since exposure to 32°C (day 8, end of experiment). On day 4, the photochemical efficiency of WT10- and Native-corals was lower than their counterparts under ambient temperature (beta regression, $p = 0.008$, $p < 0.001$, respectively). This pattern remained consistent on day 8 (beta regression, $p < 0.001$ for both) (Table S6). SS8- and HOM-corals showed no difference in photochemical efficiency between ambient and elevated temperatures on all measurement days. * indicates statistical significance between ambient and elevated temperatures within that coral group.

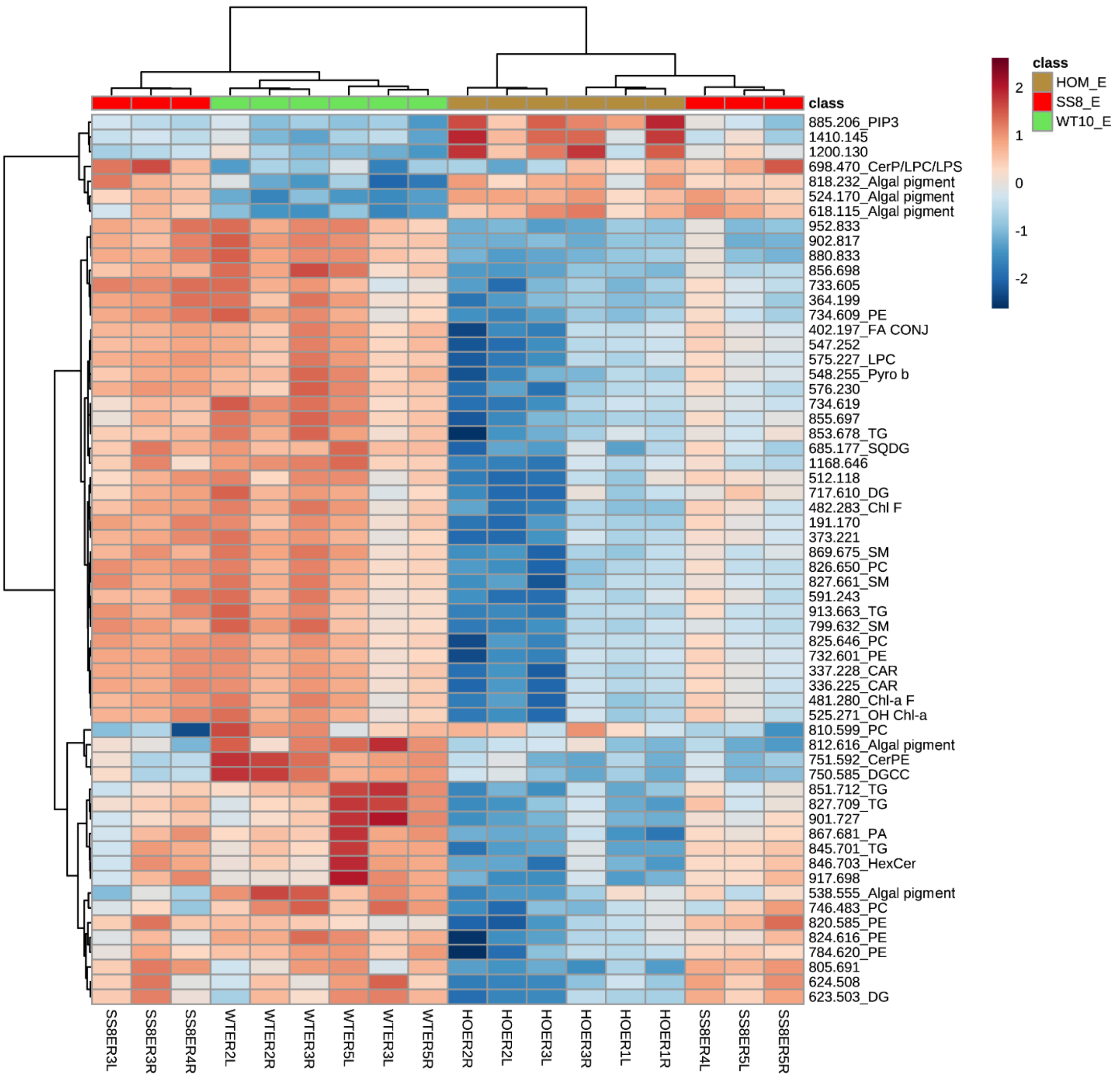

Fig. S4. Heatmap of the top 60 metabolites with a significant intensity difference between SS8-, WT10- and HOM-corals, under elevated temperature. The color scale indicates log2 fold change relative to the mean. E = elevated temperature.

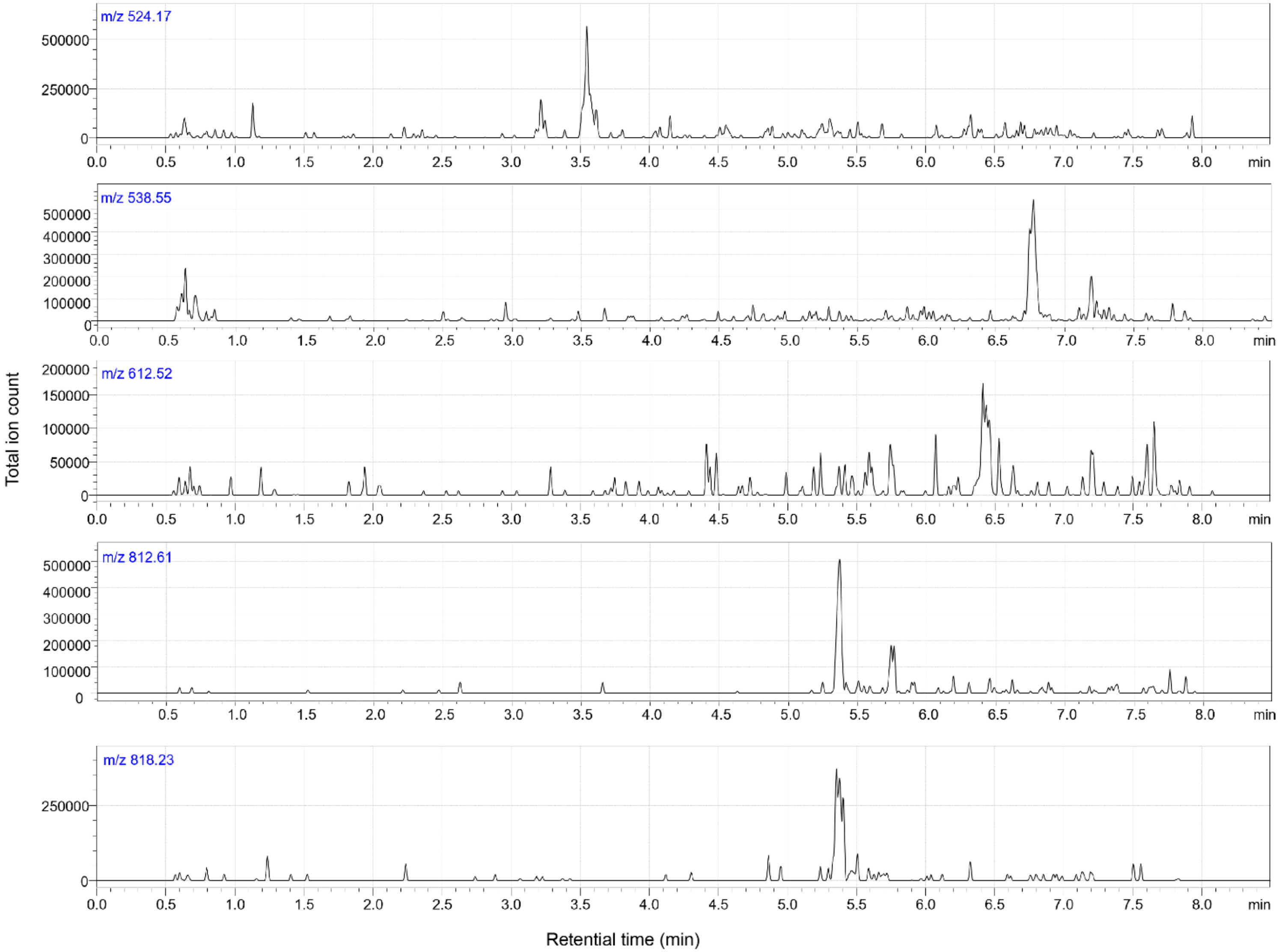

Fig. S5. Ion chromatograms for masses corresponding to algal pigment detected in both MALDI-MSI and HPLC-UV/MS on pigments extracted from coral tissues.

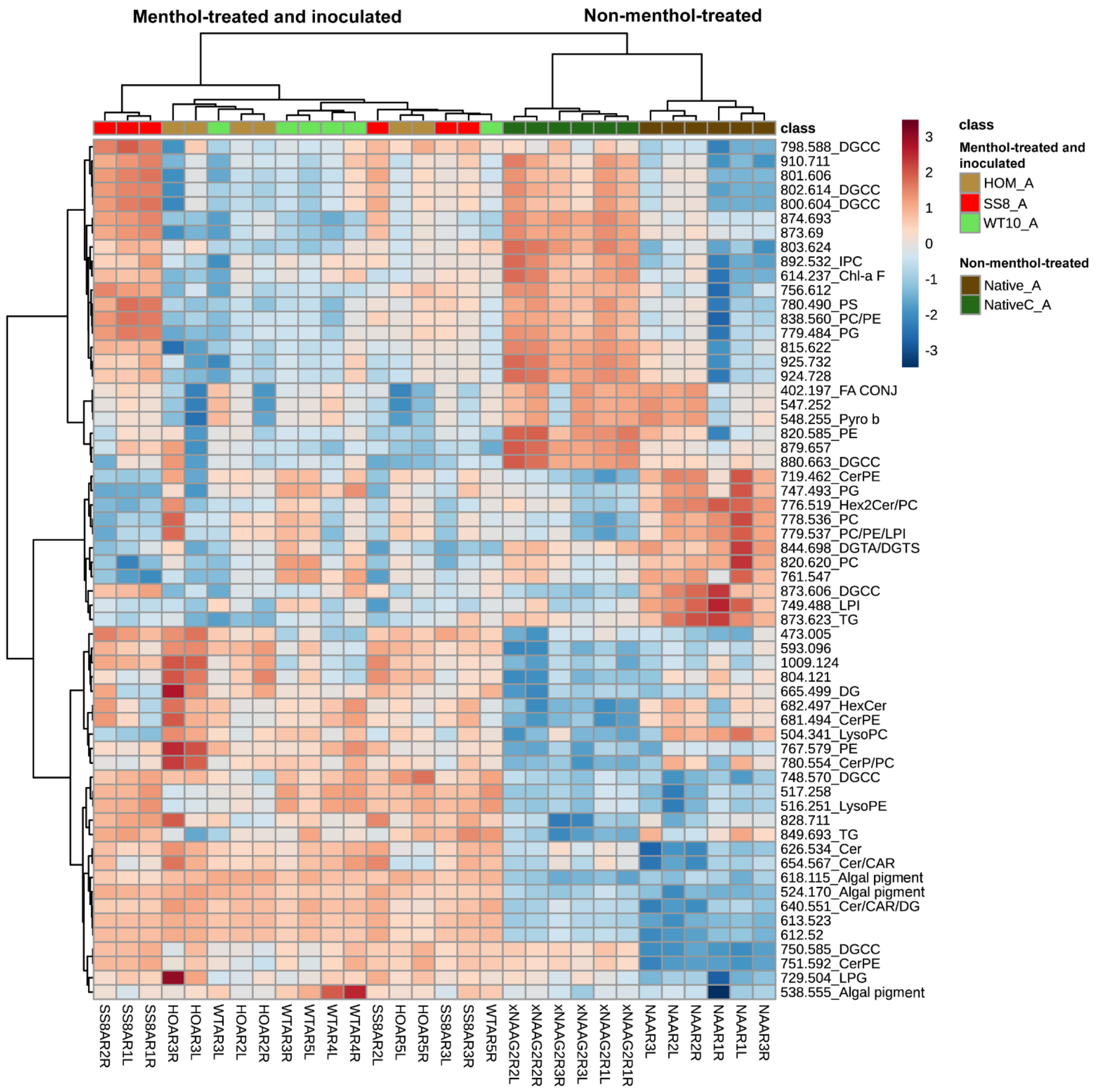

Fig. S6. Heatmap of the top 60 metabolites that were significantly different in relative intensities among coral groups, under ambient temperature. Menthol-treated and inoculated groups are SS8-, WT10-, HOM-corals; whereas control groups (non-menthol-treated) are Native- and NativeC-corals. The color scale indicates log2 fold change relative to the mean. A = ambient temperature.

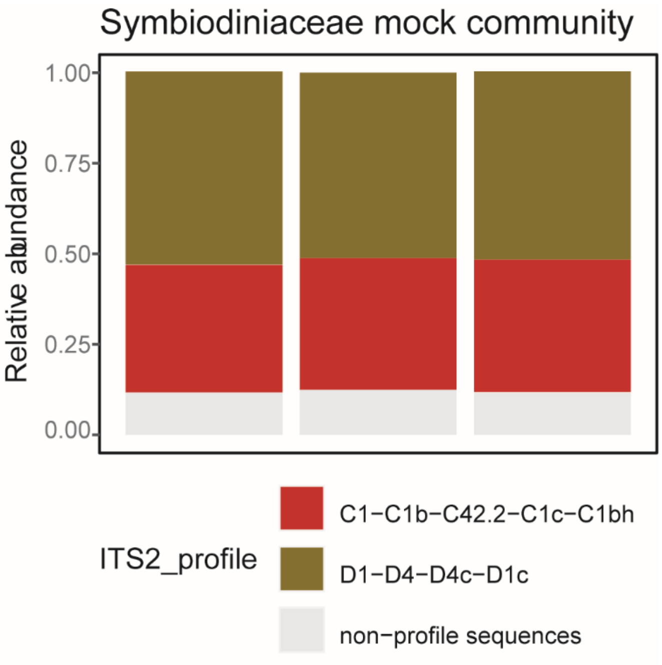

Fig. S7. ITS2 profile of the mock community samples across the three sequencing runs of this study. DNA from the mock community cultures was extracted and amplified separately; their amplicons were quantified with PicoGreen and an equal amount from each culture was pooled to make three 50:50 *Cladocopium*: *Durusdinium* mock communities.

Table S1. Experimental timeline and sampling details.

| Actual start date | Week | Event | Sampling | Replicate number |
|---|---|---|---|---|
| 29/10/2020 | -9 | Coral collection | NA | NA |
| 20/11/2020 | -6 | Coral arrival and fragmentation | Symbiodiniacea community | 1 replicate x 5 coral genotypes, *n* = 5 |
| 20/11/2020 | -6 to -5 | Coral recovery | NA | NA |
| 01/12/2020 | -4 to -1 | Menthol bleaching | NA | NA |
| 29/12/2020 | -1 | Menthol bleaching completed | Symbiodiniacea community<br>cell density<br>mass spectrometry imaging<br>pigment annotation | *n* = 4 (chemically bleached corals)<br>5 replicates x 4 genotypes x 2 treatments (with or without menthol) *n* = 40<br>*n* = 3 (chemically bleached corals for host metabolite annotation)<br>*n* = 1 (chemically bleached corals, each sample contained 4 polyps) |
| 31/12/2020 | 0 | Inoculation 1 | Symbiodiniacea community | 3 replicates x 3 inocula, *n* = 9 |
| 14/01/2021 | 4 | Inoculation 2 | NA | NA |
| 19/02/2021 | 7 | Inoculation 3 | Symbiodiniacea community | 4 replicates x 4 tanks x 4 coral groups + 8 negative control, *n* = 72 |
| 04/05/2021 | 18 | Day 0 sampling, start of heat shock | Symbiodiniacea community<br>cell density<br>coral growth<br>photochemical efficiency | 6 replicates x 4 coral groups x 1 temperature, *n* = 24<br>6 replicates x 4 coral groups x 1 temperature, *n* = 24<br>all corals, *n* = 120<br>all corals, *n* = 120 |
| 11/05/2021 | 19 | Day 4 | photochemical efficiency | all corals, *n* = 120 |

| | | | | |
|---|---|---|---|---|
| 15/05/2021 | 20 | Day 8 sampling, end of heat shock | Symbiodiniacea community<br>cell density<br>coral survival<br>photochemical efficiency<br>mass spectrometry imaging<br><br>pigment annotation | 6 replicates x 4 coral groups x 2 temperatures, *n* = 48<br>6 replicates x 4 coral groups x 2 temperatures, *n* = 48<br>all corals, *n* = 120<br>all corals, *n* = 120<br>3 biological replicates x 2 technical replicates x 5 group groups x 2 temperatures, *n* = 60<br>*n* = 2 (WT10-corals and HOM-corals, each sample contained 4 polyps) |
| 17/02/2022 | 52 | year 1 | Symbiodiniacea community | *n* = 4 |
| 13/02/2023 | 104 | year 2 | Symbiodiniacea community | *n* = 4 |

Table S2. The number of coral fragments and mortality post menthol bleaching of the *Galaxea fascicularis* colonies.

| Coral genotype | No. of fragments | No. of surviving fragments post menthol bleaching | Menthol bleaching mortality (%) |
|---|---|---|---|
| G1 | 50 | 48 | 4.0 |
| G2 | 40 | 16 | 60.0 |
| G3 | 50 | 48 | 4.0 |
| G4 | 45 | 45 | 0.0 |
| G5 | 48 | 46 | 4.2 |
| Total | 233 | 203 | 12.9 |

Table S3. Diurnal light profile of the aquarium lights (Zetlight ZP3600) during the inoculation and heat stress experiment.

| Light cycle | Duration | Time | Colour intensity setting: white, blue, colour, violet | Light intensity at coral level ($\mu E\ m^{-2}\ s^{-1}$) |
|---|---|---|---|---|
| Twilight | 0130 | 07:00-08:30 | 5, 5, 0, 5 | 8-13 |
| Sunrise | 0200 | 08:30-10:30 | 12, 18, 6, 12 | 45-55 |
| Full sun | 0500 | 10:30-15:30 | 16, 21, 11, 16 | 80-85 |
| Sunset | 0200 | 15:30-17:30 | 12, 18, 6, 12 | 45-55 |
| Twilight | 0130 | 17:30-19:00 | 10, 15, 5, 10 | 8-13 |

Table S4. Measured experimental seawater conditions. Values are mean ± 2 standard deviation.

| Parameter | Description and unit | Ambient | Elevated |
|---|---|---|---|
| Temperature | (°C) | 27.0 ± 1.0 | 32.0 ± 1.6 |
| $pH_T$ | pH in total scale | 8.2 ± 0 | 8.2 ± 0 |
| $A_T$ | Total alkalinity ($\mu mol\ kg^{-1}$) | 2320 ± 114 | 2290 ±135 |
| $\Omega_{arag}$ | Aragonite saturation state | 4.8 ± 0.3 | 5.4 ± 0.3 |
| $pCO_2^-$ | Partial pressure of $CO_2$ of air in equilibrium with seawater (µatm) | 247 ±13 | 235 ±15 |
| DIC | Dissolved inorganic carbon ($\mu mol\ kg^{-1}$) | 1885 ± 99 | 1808 ±115 |
| $HCO_3^-$ | Bicarbonate ion concentration ($\mu mol\ kg^{-1}$) | 1579 ± 83 | 1474 ±94 |
| $CO_3^{2-}$ | Carbonate ion concentration ($\mu mol\ kg^{-1}$) | 300 ± 16 | 329 ± 21 |
| Salinity | (ppt) | 35 ± 0 | 35 ± 0 |

Table S5. Pairwise comparison results (Dunn's test) of Symbiodiniaceae cell densities of coral genotypes (G1, G3, G4, G5) prior to and after four weeks of menthol treatment. P-values were corrected with the Benjamini-Hochberg method.

| Menthol treatment | Comparison | Z | $P_{adj}$ | Significance |
|---|---|---|---|---|
| No | G1 vs G3 | 2.62 | 0.053 | * |
| No | G1 vs G4 | 2.41 | 0.048 | * |
| No | G3 vs G4 | -0.21 | 0.831 | |
| No | G1 vs G5 | 0.43 | 0.803 | |
| No | G3 vs G5 | -2.19 | 0.057 | |
| No | G4 vs G5 | -1.98 | 0.072 | |
| Yes | G1 vs G3 | 3.40 | 0.004 | ** |
| Yes | G1 vs G4 | 2.54 | 0.033 | * |
| Yes | G3 vs G4 | -0.86 | 0.471 | |
| Yes | G1 vs G5 | 2.09 | 0.074 | |
| Yes | G3 vs G5 | -1.31 | 0.285 | |
| Yes | G4 vs G5 | -0.45 | 0.649 | |

Table S6. Results of negative binomial generalized liner model on coral Symbiodiniaceae cell densities of experimental groups under ambient and elevated temperatures, 20 weeks after the first inoculation.

| Temperature | Coral group | | | Estimate | SE | z value | p value | Significance |
|---|---|---|---|---|---|---|---|---|
| Ambient vs Elevated | Native-corals | | | -0.12 | 0.06 | -2.13 | 0.033 | * |
| Ambient vs Elevated | HOM-corals | | | 0.03 | 0.06 | 0.53 | 0.596 | |
| Ambient vs Elevated | SS8-corals | | | -0.01 | 0.06 | -0.21 | 0.830 | |
| Ambient vs Elevated | WT10-corals | | | -2.20 | 0.06 | -39.14 | < 0.001 | *** |
| Ambient | Native-corals | vs | HOM-corals | -0.42 | 0.06 | -7.44 | < 0.001 | *** |
| Ambient | Native-corals | vs | SS8-corals | -0.18 | 0.06 | -3.30 | < 0.001 | *** |
| Ambient | Native-corals | vs | WT10-corals | -0.20 | 0.06 | -3.51 | < 0.001 | *** |
| Ambient | HOM-corals | vs | SS8-corals | 0.23 | 0.06 | 4.14 | < 0.001 | *** |
| Ambient | HOM-corals | vs | WT10-coral | 0.22 | 0.06 | 3.92 | < 0.001 | *** |
| Ambient | SS8-corals | vs | WT10-corals | -0.01 | 0.06 | -0.21 | 0.830 | |
| Elevated | WT10-corals | vs | SS8-corals | 2.20 | 0.06 | 39.13 | < 0.001 | *** |
| Elevated | WT10-corals | vs | HOM-corals | 2.01 | 0.06 | 35.74 | < 0.001 | *** |
| Elevated | WT10-corals | vs | Native-corals | 2.27 | 0.06 | 40.52 | < 0.001 | *** |
| Elevated | HOM-corals | vs | Native-corals | 0.27 | 0.06 | 4.78 | < 0.001 | *** |
| Elevated | SS8-corals | vs | HOM-corals | -0.19 | 0.06 | -3.39 | 0.001 | *** |
| Elevated | SS8-corals | vs | Native-corals | 0.08 | 0.06 | 1.38 | 0.167 | |

Table S7. Pairwise comparison results (TukeyHSD) of coral size 18 weeks after the first inoculation. The corals were maintained at ambient temperature during this time.

| Comparison | | | Diff | Lower | Upper | $P_{adj}$ | Significance |
|---|---|---|---|---|---|---|---|
| WT10-corals | vs | SS8-corals | -0.85 | -4.51 | 2.80 | 0.930 | |
| HOM-corals | vs | SS8-corals | -3.22 | -6.01 | -0.43 | 0.017 | * |
| Native-corals | vs | SS8-corals | -5.03 | -7.84 | -2.23 | < 0.001 | *** |
| HOM-corals | vs | WT10-corals | -2.37 | -5.73 | 1.00 | 0.262 | |
| Native-corals | vs | WT10-corals | -4.18 | -7.56 | -0.81 | 0.009 | ** |
| Native-corals | vs | HOM-corals | -1.81 | -4.23 | 0.60 | 0.211 | |

Table S8. Pairwise comparison results (Dunn's test) of coral survival under elevated temperature, 20 weeks after the first inoculation. P-values were corrected with the Benjamini-Hochberg method.

| Comparison | | | Z | $P_{adj}$ | Significance |
|---|---|---|---|---|---|
| HOM-corals | vs | Native-corals | -1.75 | 0.096 | |
| HOM-corals | vs | SS8-corals | -2.05 | 0.060 | |
| Native-corals | vs | SS8-corals | -0.53 | 0.592 | |
| HOM-corals | vs | WT10-corals | 2.67 | 0.015 | * |
| Native-corals | vs | WT10-corals | 3.93 | < 0.001 | *** |
| SS8-corals | vs | WT10-corals | 4.04 | < 0.001 | *** |

Table S9. Results of beta regressions on photochemical efficiency (dark-adapted maximum photosystem II quantum yield, Fv/Fm) of the different experimental groups under ambient and elevated temperatures, 20 weeks after the first inoculation.

| Temperature | Coral group | | | Estimate | SE | z value | p value | Significance |
|---|---|---|---|---|---|---|---|---|
| Ambient vs Elevated | Native-corals | | | -0.12 | 0.03 | -4.07 | < 0.001 | *** |
| Ambient vs Elevated | HOM-corals | | | 0.05 | 0.03 | 1.86 | 0.063 | |
| Ambient vs Elevated | SS8-corals | | | -0.02 | 0.04 | -0.59 | 0.555 | |
| Ambient vs Elevated | WT10-corals | | | -1.04 | 0.06 | -18.65 | < 0.001 | *** |
| Ambient | Native-corals | vs | HOM-corals | -0.26 | 0.03 | -8.83 | < 0.001 | *** |
| Ambient | Native-corals | vs | SS8-corals | -0.21 | 0.03 | -6.45 | < 0.001 | *** |
| Ambient | Native-corals | vs | WT10-corals | -0.18 | 0.04 | -4.28 | < 0.001 | *** |
| Ambient | HOM-corals | vs | SS8-corals | 0.05 | 0.03 | 1.46 | 0.144 | |
| Ambient | HOM-corals | vs | WT10-corals | 0.08 | 0.04 | 1.99 | 0.047 | * |
| Ambient | SS8-corals | vs | WT10-corals | 0.03 | 0.04 | 0.78 | 0.433 | |
| Elevated | WT10-corals | vs | SS8-corals | 0.98 | 0.05 | 19.03 | < 0.001 | *** |
| Elevated | WT10-corals | vs | HOM-corals | 1.01 | 0.05 | 21.43 | < 0.001 | *** |
| Elevated | WT10-corals | vs | Native-corals | 1.10 | 0.05 | 23.23 | < 0.001 | *** |
| Elevated | HOM-corals | vs | Native-corals | 0.09 | 0.03 | 2.92 | 0.004 | ** |
| Elevated | SS8-corals | vs | HOM-corals | 0.03 | 0.04 | 0.82 | 0.412 | |
| Elevated | SS8-corals | vs | Native-corals | 0.12 | 0.04 | 3.20 | 0.001 | ** |

Table S10. Abbreviations and details of annotated metabolites.

| Abbreviation | Full name | Category |
|---|---|---|
| CAR | Acyl carnitine | Energy homeostasis |
| Car | Carotenoid | Photosynthesis/oxidative stress |
| Cer | Ceramide | Signaling |
| CerPE | Ceramide phosphoethanolamine | Signaling |
| CerP | Ceramide-phosphate | Signaling/sphingosine rheostat |
| Chl a | Chlorophyll a | Energy/photosynthesis/oxidative stress |
| Chl-a F | Chlorophyll a fragment | Energy/photosynthesis/oxidative stress |
| Chl F | Chlorophyll fragment | Energy/photosynthesis |
| DG | Diglyceride | Energy |
| DGCC | Diacylglycerylcarboxyhydroxymethylcholine | Structural |
| DGDG | Digalactosyldiacylglycerol | Structural |
| DGTA | Diacylglycerylhydroxymethyltrimethylalanine | Structural |
| DGTS | Diacylglyceryltrimethylhomoserine | Structural |
| FA | Fatty acids/esters | Energy/backbone |
| FA CONJ | Fatty acid conjugate | Energy/signaling |
| GlcCer | Glucosylceramide | Signaling/immune/cellular recognition |
| HexCer | Hexosylceramide | Signaling/immune/cellular recognition |
| HG | Headgroup | Structural |
| IPC | Inositolphosphoryl-ceramides | Signaling/apoptosis |
| LPA | Lysophosphatidic acid | Signaling |
| LPC | Lysophosphatidylcholine | Structural |
| LPI | Lysophosphatidylinositol | Structural |
| LPG | Lysophosphatidylglycerol | Structural |
| MG | Monoacylglycerol | Energy |
| MGCC | Monoacylglycerylcarboxyhydroxymethylcholine | Structural |
| MGDG | Monogalactosyldiacylglycerol | Photosynthesis |
| MIPC | Mannosylinositol phosphorylceramide | Signaling/immune/cellular recognition |
| PA | Phosphatidic acid | Signaling |
| PC | Glycerophosphocholine | Structural |
| PE | Phosphatidylethanolamine | Structural |
| PG | Phosphoglycan | Structural/immune |
| Phe a | Pheophytin a | Photosynthesis/oxidative stress |
| Pheo a | Pheophorbide a | Energy/photosynthesis |
| PI | Phosphatidylinositol | Structural/signaling |
| PIP | Phosphatidylinositol-*n*-phosphate | Structural/signaling |
| PM | Polar metabolite | All |
| Pyro b | Pyropheophorbide b | Photosynthesis/oxidative stress |
| PS | Phosphatidylserine | Structural/signaling |

| | | |
|---|---|---|
| SQDG | Sulfoquinovosyl diacylglycerol | Structural/energy/photosynthesis |
| ST | Sterol | Structural/signaling |
| TG | Triglyceride | Energy |

Table S11. Metabolites with significant differences in relative intensities between SS8-, WT10, HOM-corals under elevated temperature based on an ANOVA. E = elevated temperature.

| m/z | Annotation | F.value | FDR | Tukey's HSD |
|---|---|---|---|---|
| 191.17 | | 22.8 | 0.001 | SS8_E-HOM_E; WT10_E-HOM_E |
| 301.091 | | 10.5 | 0.006 | SS8_E-HOM_E; WT10_E-HOM_E |
| 302.095 | | 11.5 | 0.005 | SS8_E-HOM_E; WT10_E-HOM_E |
| 336.225 | CAR | 16.2 | 0.002 | SS8_E-HOM_E; WT10_E-HOM_E |
| 337.228 | CAR | 19.1 | 0.001 | SS8_E-HOM_E; WT10_E-HOM_E |
| 364.199 | | 17.6 | 0.002 | SS8_E-HOM_E; WT10_E-HOM_E |
| 373.221 | | 16.0 | 0.002 | SS8_E-HOM_E; WT10_E-HOM_E |
| 402.197 | FA CONJ | 16.7 | 0.002 | SS8_E-HOM_E; WT10_E-HOM_E |
| 464.3485 | CAR | 7.5 | 0.018 | SS8_E-HOM_E; WT10_E-HOM_E |
| 472.362 | DGTS | 10.6 | 0.006 | SS8_E-HOM_E; WT10_E-HOM_E |
| 478.328 | LPE/CAR | 11.9 | 0.005 | SS8_E-HOM_E; WT10_E-HOM_E |
| 481.28 | Chl-a F | 19.9 | 0.001 | SS8_E-HOM_E; WT10_E-HOM_E |
| 482.283 | Chl F | 29.4 | < 0.001 | SS8_E-HOM_E; WT10_E-HOM_E |
| 482.359 | PC/LPE | 5.9 | 0.034 | WT10_E-SS8_E |
| 490.137 | | 11.5 | 0.005 | SS8_E-HOM_E; WT10_E-HOM_E |
| 491.139 | | 12.1 | 0.004 | SS8_E-HOM_E; WT10_E-HOM_E |
| 492.38 | LPC | 7.5 | 0.018 | SS8_E-HOM_E; WT10_E-HOM_E |
| 504.341 | LPC | 9.6 | 0.009 | SS8_E-HOM_E; WT10_E-SS8_E |
| 506.36 | CerP/LPC | 8.2 | 0.014 | SS8_E-HOM_E; WT10_E-HOM_E |
| 511.158 | | 7.5 | 0.018 | SS8_E-HOM_E; WT10_E-HOM_E |
| 512.118 | | 15.9 | 0.002 | SS8_E-HOM_E; WT10_E-HOM_E |
| 516.251 | LPE | 6.2 | 0.030 | SS8_E-HOM_E; WT10_E-SS8_E |
| 524.17 | Algal pigment | 132.0 | < 0.001 | WT10_E-HOM_E; WT10_E-SS8_E |
| 525.271 | OH Chl-a | 17.7 | 0.002 | SS8_E-HOM_E; WT10_E-HOM_E |
| 538.555 | Algal pigment (Car) | 18.2 | 0.001 | WT10_E-HOM_E; WT10_E-SS8_E |
| 544.362 | LPC/Car | 9.4 | 0.009 | SS8_E-HOM_E; WT10_E-HOM_E |
| 547.252 | | 19.7 | 0.001 | SS8_E-HOM_E; WT10_E-HOM_E |
| 548.255 | Pyro b | 26.7 | < 0.001 | SS8_E-HOM_E; WT10_E-HOM_E |
| 554.36 | | 6.6 | 0.026 | WT10_E-HOM_E |
| 562.373 | MGCC | 7.5 | 0.019 | SS8_E-HOM_E; WT10_E-HOM_E |
| 575.227 | LPC | 17.3 | 0.002 | SS8_E-HOM_E; WT10_E-HOM_E |
| 576.23 | | 17.3 | 0.002 | SS8_E-HOM_E; WT10_E-HOM_E |
| 591.243 | | 14.2 | 0.002 | SS8_E-HOM_E; WT10_E-HOM_E |
| 618.115 | Algal pigment | 51.0 | < 0.001 | WT10_E-HOM_E; WT10_E-SS8_E |
| 620.221 | | 12.0 | 0.005 | SS8_E-HOM_E; WT10_E-HOM_E |
| 623.503 | DG | 17.8 | 0.002 | SS8_E-HOM_E; WT10_E-HOM_E |

| | | | | |
|---|---|---|---|---|
| 624.508 | | 19.3 | 0.001 | SS8_E-HOM_E; WT10_E-HOM_E |
| 654.567 | Cer | 7.4 | 0.019 | WT10_E-HOM_E; WT10_E-SS8_E |
| 655.566 | DG | 5.3 | 0.044 | WT10_E-HOM_E |
| 663.194 | | 14.0 | 0.003 | SS8_E-HOM_E; WT10_E-HOM_E |
| 673.176 | PI | 11.3 | 0.005 | SS8_E-HOM_E; WT10_E-HOM_E |
| 674.183 | | 11.3 | 0.005 | SS8_E-HOM_E; WT10_E-HOM_E |
| 685.177 | SQDG | 17.3 | 0.002 | SS8_E-HOM_E; WT10_E-HOM_E |
| 686.193 | | 10.7 | 0.006 | SS8_E-HOM_E; WT10_E-HOM_E |
| 689.155 | | 10.0 | 0.008 | SS8_E-HOM_E; WT10_E-HOM_E |
| 697.468 | CerPE | 9.6 | 0.008 | SS8_E-HOM_E; WT10_E-HOM_E |
| 698.47 | CerP/LPC/LPS | 15.4 | 0.002 | SS8_E-HOM_E; WT10_E-SS8_E |
| 701.158 | | 5.6 | 0.040 | WT10_E-HOM_E |
| 702.166 | | 9.9 | 0.008 | SS8_E-HOM_E; WT10_E-HOM_E |
| 705.484 | PA | 5.1 | 0.047 | WT10_E-HOM_E |
| 716.607 | HexCer/Cer | 8.6 | 0.012 | SS8_E-HOM_E; WT10_E-HOM_E |
| 717.61 | DG | 14.4 | 0.002 | SS8_E-HOM_E; WT10_E-HOM_E |
| 719.462 | CerPE | 5.1 | 0.047 | SS8_E-HOM_E; WT10_E-HOM_E |
| 732.601 | PE | 14.5 | 0.002 | SS8_E-HOM_E; WT10_E-HOM_E |
| 733.511 | DG | 5.0 | 0.050 | WT10_E-HOM_E |
| 733.605 | | 14.4 | 0.002 | SS8_E-HOM_E; WT10_E-HOM_E |
| 734.517 | PC | 6.1 | 0.032 | WT10_E-HOM_E |
| 734.609 | PE | 17.1 | 0.002 | SS8_E-HOM_E; WT10_E-HOM_E |
| 734.619 | | 20.3 | 0.001 | SS8_E-HOM_E; WT10_E-HOM_E |
| 736.284 | | 13.4 | 0.003 | SS8_E-HOM_E; WT10_E-HOM_E |
| 737.286 | PIP (fragment) | 10.8 | 0.006 | SS8_E-HOM_E; WT10_E-HOM_E |
| 742.574 | CerP/LPC/PC | 5.6 | 0.040 | WT10_E-HOM_E; WT10_E-SS8_E |
| 745.477 | PG | 9.1 | 0.010 | SS8_E-HOM_E; WT10_E-HOM_E |
| 746.483 | PC | 16.6 | 0.002 | SS8_E-HOM_E; WT10_E-HOM_E; WT10_E-SS8_E |
| 747.493 | PG | 7.0 | 0.022 | SS8_E-HOM_E; WT10_E-HOM_E |
| 748.57 | DGCC | 5.9 | 0.034 | WT10_E-SS8_E |
| 748.595 | PE | 11.2 | 0.005 | SS8_E-HOM_E; WT10_E-HOM_E |
| 749.601 | DG | 9.1 | 0.010 | SS8_E-HOM_E; WT10_E-HOM_E |
| 750.585 | DGCC | 33.8 | < 0.001 | WT10_E-HOM_E; WT10_E-SS8_E |
| 751.592 | CerPE | 38.1 | < 0.001 | WT10_E-HOM_E; WT10_E-SS8_E |
| 752.26 | | 10.6 | 0.006 | SS8_E-HOM_E; WT10_E-HOM_E |
| 754.584 | PE | 7.1 | 0.021 | SS8_E-HOM_E; WT10_E-HOM_E |
| 757.615 | DG | 5.2 | 0.046 | SS8_E-HOM_E |
| 761.547 | | 12.8 | 0.004 | SS8_E-HOM_E; WT10_E-HOM_E |
| 762.482 | PC | 11.4 | 0.005 | SS8_E-HOM_E; WT10_E-SS8_E |
| 765.52 | DGCC | 5.1 | 0.048 | SS8_E-HOM_E |

| | | | | |
|---|---|---|---|---|
| 766.575 | PC | 6.4 | 0.028 | WT10_E-HOM_E |
| 769.478 | MGDG | 5.4 | 0.043 | SS8_E-HOM_E |
| 771.61 | | 6.8 | 0.024 | WT10_E-HOM_E; WT10_E-SS8_E |
| 772.572 | DGCC | 8.4 | 0.013 | SS8_E-HOM_E; WT10_E-HOM_E |
| 775.523 | CerPE | 5.5 | 0.042 | WT10_E-HOM_E |
| 779.484 | PG | 6.6 | 0.026 | SS8_E-HOM_E |
| 780.49 | PS | 7.6 | 0.018 | SS8_E-HOM_E |
| 781.595 | PG | 6.3 | 0.028 | SS8_E-HOM_E |
| 782.606 | PE | 11.3 | 0.005 | SS8_E-HOM_E; WT10_E-HOM_E |
| 783.609 | DG | 9.8 | 0.008 | SS8_E-HOM_E; WT10_E-HOM_E |
| 784.585 | PC | 6.2 | 0.030 | WT10_E-HOM_E |
| 784.61 | DGTA/DGTS | 10.2 | 0.007 | SS8_E-HOM_E; WT10_E-HOM_E |
| 784.62 | PE | 18.1 | 0.001 | SS8_E-HOM_E; WT10_E-HOM_E |
| 788.604 | PC | 7.4 | 0.019 | SS8_E-HOM_E; WT10_E-HOM_E |
| 790.513 | | 11.8 | 0.005 | SS8_E-HOM_E; WT10_E-HOM_E |
| 796.62 | PC | 8.8 | 0.011 | SS8_E-HOM_E; WT10_E-SS8_E |
| 797.624 | | 5.1 | 0.048 | WT10_E-HOM_E |
| 798.624 | | 9.4 | 0.009 | SS8_E-HOM_E; WT10_E-HOM_E |
| 799.632 | SM | 14.2 | 0.002 | SS8_E-HOM_E; WT10_E-HOM_E |
| 803.624 | | 13.4 | 0.003 | WT10_E-HOM_E; WT10_E-SS8_E |
| 804.121 | | 8.0 | 0.015 | SS8_E-HOM_E; WT10_E-HOM_E |
| 805.691 | | 40.2 | < 0.001 | SS8_E-HOM_E; WT10_E-HOM_E |
| 806.567 | PC | 5.2 | 0.046 | WT10_E-HOM_E |
| 808.621 | PE | 6.2 | 0.030 | SS8_E-HOM_E; WT10_E-HOM_E |
| 810.599 | PC | 14.2 | 0.002 | SS8_E-HOM_E; WT10_E-SS8_E |
| 810.636 | | 9.9 | 0.008 | SS8_E-HOM_E; WT10_E-HOM_E |
| 811.606 | PC | 6.3 | 0.029 | WT10_E-SS8_E |
| 811.641 | DG/TG | 10.0 | 0.008 | SS8_E-HOM_E; WT10_E-HOM_E |
| 812.616 | Algal pigment | 21.3 | 0.001 | WT10_E-HOM_E; WT10_E-SS8_E |
| 814.62 | DGCC | 6.8 | 0.024 | SS8_E-HOM_E |
| 815.622 | | 6.2 | 0.030 | SS8_E-HOM_E |
| 816.628 | PE | 11.5 | 0.005 | SS8_E-HOM_E; WT10_E-HOM_E |
| 818.232 | Algal pigment | 23.8 | 0.001 | WT10_E-HOM_E; WT10_E-SS8_E |
| 820.585 | PE | 17.9 | 0.002 | SS8_E-HOM_E; WT10_E-HOM_E |
| 820.62 | PC | 7.0 | 0.022 | WT10_E-HOM_E |
| 821.623 | | 5.7 | 0.038 | WT10_E-HOM_E |
| 823.63 | PC | 8.7 | 0.012 | SS8_E-HOM_E; WT10_E-HOM_E |
| 824.616 | PE | 18.2 | 0.001 | SS8_E-HOM_E; WT10_E-HOM_E |
| 824.652 | | 11.6 | 0.005 | SS8_E-HOM_E; WT10_E-SS8_E |
| 825.646 | PC | 15.5 | 0.002 | SS8_E-HOM_E; WT10_E-HOM_E |

| | | | | |
|---|---|---|---|---|
| 826.65 | PC | 17.2 | 0.002 | SS8_E-HOM_E; WT10_E-HOM_E |
| 827.661 | SM | 16.2 | 0.002 | SS8_E-HOM_E; WT10_E-HOM_E |
| 827.709 | TG | 18.7 | 0.001 | SS8_E-HOM_E; WT10_E-HOM_E |
| 828.633 | DGCC | 7.3 | 0.020 | WT10_E-HOM_E; WT10_E-SS8_E |
| 828.666 | | 7.1 | 0.021 | SS8_E-HOM_E; WT10_E-HOM_E |
| 829.728 | DG/TG | 13.7 | 0.003 | SS8_E-HOM_E; WT10_E-HOM_E |
| 834.641 | | 7.2 | 0.020 | SS8_E-HOM_E; WT10_E-HOM_E |
| 835.599 | | 5.3 | 0.044 | WT10_E-HOM_E |
| 838.56 | PC/PE | 5.6 | 0.038 | WT10_E-SS8_E |
| 838.634 | PC | 8.1 | 0.014 | SS8_E-HOM_E; WT10_E-SS8_E |
| 843.691 | MGDG | 12.7 | 0.004 | SS8_E-HOM_E; WT10_E-HOM_E |
| 844.698 | DGTA/DGTS | 7.8 | 0.016 | SS8_E-HOM_E; WT10_E-HOM_E |
| 845.701 | TG | 22.3 | 0.001 | SS8_E-HOM_E; WT10_E-HOM_E |
| 846.703 | HexCer | 18.7 | 0.001 | SS8_E-HOM_E; WT10_E-HOM_E |
| 849.693 | TG | 10.0 | 0.008 | WT10_E-HOM_E |
| 850.616 | DGCC | 7.6 | 0.018 | SS8_E-HOM_E; WT10_E-HOM_E |
| 851.712 | TG | 31.5 | < 0.001 | SS8_E-HOM_E; WT10_E-HOM_E; WT10_E-SS8_E |
| 852.646 | | 9.0 | 0.010 | SS8_E-HOM_E; WT10_E-HOM_E |
| 853.678 | TG | 16.7 | 0.002 | SS8_E-HOM_E; WT10_E-HOM_E |
| 855.659 | TG | 8.2 | 0.014 | SS8_E-HOM_E; WT10_E-HOM_E |
| 855.697 | | 28.4 | < 0.001 | SS8_E-HOM_E; WT10_E-HOM_E; WT10_E-SS8_E |
| 856.698 | | 28.4 | < 0.001 | SS8_E-HOM_E; WT10_E-HOM_E; WT10_E-SS8_E |
| 857.675 | CerPE/TG | 11.0 | 0.006 | SS8_E-HOM_E; WT10_E-HOM_E |
| 867.681 | PA | 27.2 | < 0.001 | SS8_E-HOM_E; WT10_E-HOM_E |
| 869.675 | SM | 20.3 | 0.001 | SS8_E-HOM_E; WT10_E-HOM_E |
| 871.718 | TG | 8.9 | 0.011 | SS8_E-HOM_E; WT10_E-HOM_E |
| 873.623 | TG | 6.5 | 0.026 | SS8_E-HOM_E; WT10_E-HOM_E |
| 879.657 | | 5.9 | 0.035 | SS8_E-HOM_E |
| 880.663 | DGCC | 8.9 | 0.011 | SS8_E-HOM_E; WT10_E-HOM_E |
| 880.833 | | 15.4 | 0.002 | SS8_E-HOM_E; WT10_E-HOM_E |
| 881.676 | | 6.5 | 0.026 | SS8_E-HOM_E |
| 883.69 | PG/CerPE | 6.4 | 0.028 | WT10_E-HOM_E |
| 884.695 | DGCC | 6.1 | 0.031 | WT10_E-HOM_E; WT10_E-SS8_E |
| 885.206 | PIP3 | 42.9 | < 0.001 | SS8_E-HOM_E; WT10_E-HOM_E |
| 899.683 | | 11.7 | 0.005 | SS8_E-HOM_E; WT10_E-HOM_E |
| 900.69 | HexCer | 12.4 | 0.004 | SS8_E-HOM_E; WT10_E-HOM_E |
| 901.727 | | 16.3 | 0.002 | SS8_E-HOM_E; WT10_E-HOM_E |
| 902.817 | | 15.4 | 0.002 | SS8_E-HOM_E; WT10_E-HOM_E |
| 906.174 | | 5.3 | 0.045 | SS8_E-HOM_E; WT10_E-HOM_E |
| 913.663 | TG | 14.9 | 0.002 | SS8_E-HOM_E; WT10_E-HOM_E |

| | | | | |
|---|---|---|---|---|
| 914.668 | TG | 6.0 | 0.032 | SS8_E-HOM_E |
| 917.698 | | 16.0 | 0.002 | SS8_E-HOM_E; WT10_E-HOM_E |
| 924.728 | | 9.8 | 0.008 | SS8_E-HOM_E |
| 925.732 | | 6.9 | 0.023 | SS8_E-HOM_E |
| 952.833 | | 16.6 | 0.002 | SS8_E-HOM_E; WT10_E-HOM_E |
| 953.834 | TG | 10.9 | 0.006 | SS8_E-HOM_E; WT10_E-HOM_E |
| 979.608 | | 10.3 | 0.007 | SS8_E-HOM_E; WT10_E-HOM_E |
| 980.616 | | 5.5 | 0.041 | WT10_E-HOM_E |
| 981.627 | | 11.2 | 0.005 | SS8_E-HOM_E; WT10_E-HOM_E |
| 988.101 | | 5.3 | 0.045 | WT10_E-HOM_E |
| 988.683 | | 9.0 | 0.010 | SS8_E-HOM_E; WT10_E-HOM_E |
| 1007.648 | | 7.3 | 0.020 | WT10_E-HOM_E |
| 1008.648 | | 6.6 | 0.026 | WT10_E-HOM_E |
| 1009.124 | | 10.3 | 0.007 | SS8_E-HOM_E; WT10_E-HOM_E |
| 1030.717 | | 6.7 | 0.025 | WT10_E-HOM_E |
| 1032.663 | | 5.2 | 0.045 | SS8_E-HOM_E; WT10_E-HOM_E |
| 1033.662 | | 5.7 | 0.038 | SS8_E-HOM_E; WT10_E-HOM_E |
| 1073.23 | | 5.3 | 0.044 | SS8_E-HOM_E; WT10_E-HOM_E |
| 1089.209 | | 8.2 | 0.014 | SS8_E-HOM_E; WT10_E-HOM_E |
| 1090.211 | | 13.6 | 0.003 | SS8_E-HOM_E; WT10_E-HOM_E |
| 1168.646 | | 15.2 | 0.002 | SS8_E-HOM_E; WT10_E-HOM_E |
| 1200.13 | | 14.5 | 0.002 | SS8_E-HOM_E; WT10_E-HOM_E |
| 1235.638 | | 5.2 | 0.046 | SS8_E-HOM_E; WT10_E-HOM_E |
| 1410.145 | | 21.3 | 0.001 | SS8_E-HOM_E; WT10_E-HOM_E |
| 1417.621 | | 12.9 | 0.004 | SS8_E-HOM_E; WT10_E-HOM_E |
| 1418.632 | | 5.5 | 0.041 | WT10_E-HOM_E |

Table S12. T-test results of metabolites that were significantly different in relative intensities between SS8- and WT10 corals under elevated temperature. A fold change (FC) > 1 indicates that this metabolite was enriched in WT10-corals.

| m/z | Annotation | T.stat | $P_{adj}$ | FC | Log2(FC) |
|---|---|---|---|---|---|
| 524.17 | Algal pigment | 13.69 | < 0.001 | 79.32 | 6.31 |
| 538.555 | Algal pigment (Car) | -5.04 | 0.021 | 0.30 | -1.75 |
| 618.115 | Algal pigment | 8.14 | 0.002 | 11.87 | 3.57 |
| 654.567 | Cer | -4.58 | 0.035 | 0.45 | -1.16 |
| 698.47 | CerP/LPC/LPS | 6.59 | 0.005 | 3.84 | 1.94 |
| 750.585 | DGCC | -6.48 | 0.005 | 0.21 | -2.23 |
| 751.592 | CerPE | -7.17 | 0.004 | 0.17 | -2.52 |
| 803.624 | | -4.49 | 0.037 | 0.34 | -1.56 |
| 810.599 | PC | -4.90 | 0.024 | 0.71 | -0.49 |
| 812.616 | Algal pigment | -5.38 | 0.016 | 0.52 | -0.94 |
| 818.232 | Algal pigment | 5.57 | 0.014 | 3.04 | 1.60 |
| 824.652 | | -6.85 | 0.005 | 0.53 | -0.91 |
| 981.551 | | 5.05 | 0.021 | 3.06 | 1.62 |

Table S13. T-test results of metabolites that were significantly different in relative intensities between WT10-corals under ambient versus elevated temperatures. A fold change (FC) > 1 suggests that this metabolite was enriched under ambient temperature, whereas a FC < 1 indicates that the metabolite was enriched under elevated temperature. For instance, a FC value of 0.1 means that this metabolite was 10 times more abundant elevated temperature.

| m/z | Annotation | T.stat | FDR | FC | Log2(FC) |
|---|---|---|---|---|---|
| 472.362 | DGTS | -4.41 | 0.014 | 0.55 | -0.87 |
| 504.341 | LPC | -4.73 | 0.010 | 0.35 | -1.53 |
| 516.251 | LPE | 3.77 | 0.032 | 2.75 | 1.46 |
| 517.258 | | 3.85 | 0.029 | 3.18 | 1.67 |
| 524.17 | Algal pigment | 15.52 | < 0.001 | 81.27 | 6.34 |
| 612.52 | | 22.86 | < 0.001 | 8.40 | 3.07 |
| 613.523 | | 19.28 | < 0.001 | 15.93 | 3.99 |
| 618.115 | Algal pigment | 13.40 | < 0.001 | 14.45 | 3.85 |
| 626.534 | Cer | 7.40 | 0.001 | 2.35 | 1.23 |
| 640.551 | Cer/CAR/DGTS | 10.40 | < 0.001 | 3.30 | 1.72 |
| 654.567 | Cer | 4.29 | 0.017 | 1.48 | 0.56 |
| 665.499 | DG | 5.00 | 0.007 | 1.90 | 0.92 |
| 681.494 | CerPE | 5.81 | 0.002 | 1.98 | 0.98 |
| 682.497 | HexCer | 6.12 | 0.002 | 2.68 | 1.42 |
| 697.468 | CerPE | 3.68 | 0.033 | 2.59 | 1.37 |
| 698.47 | CerP/LPC/LPS | 3.99 | 0.025 | 3.22 | 1.69 |
| 745.456 | | 7.19 | 0.001 | 5.00 | 2.32 |
| 750.585 | DGCC | -6.58 | 0.002 | 0.24 | -2.04 |
| 751.592 | CerPE | -6.32 | 0.002 | 0.23 | -2.13 |
| 756.612 | | -5.28 | 0.005 | 0.45 | -1.14 |
| 757.615 | DG | -4.81 | 0.009 | 0.42 | -1.26 |
| 766.575 | PC | 8.42 | < 0.001 | 2.42 | 1.28 |
| 767.579 | PE | 8.72 | < 0.001 | 2.97 | 1.57 |
| 768.593 | PC | 3.51 | 0.039 | 1.50 | 0.58 |
| 769.594 | PC | 3.38 | 0.047 | 1.46 | 0.55 |
| 774.587 | DGCC | 4.00 | 0.025 | 2.17 | 1.12 |
| 775.591 | TG | 3.55 | 0.038 | 2.38 | 1.25 |
| 780.591 | | 6.00 | 0.002 | 2.61 | 1.38 |
| 781.595 | PG | 6.21 | 0.002 | 3.10 | 1.63 |
| 788.556 | PC | 3.95 | 0.026 | 2.15 | 1.10 |
| 789.56 | PC | 3.74 | 0.033 | 2.41 | 1.27 |
| 794.605 | | 6.34 | 0.002 | 2.26 | 1.17 |
| 795.608 | | 7.11 | 0.001 | 3.11 | 1.64 |

| | | | | | |
|---|---|---|---|---|---|
| 796.62 | PC | 3.34 | 0.048 | 1.50 | 0.59 |
| 803.624 | | -6.00 | 0.002 | 0.16 | -2.65 |
| 804.529 | | 5.81 | 0.002 | 3.76 | 1.91 |
| 805.532 | DG/TG | 5.66 | 0.003 | 5.53 | 2.47 |
| 806.547 | | 3.43 | 0.043 | 1.65 | 0.72 |
| 808.621 | PE | 4.65 | 0.010 | 2.51 | 1.33 |
| 810.599 | PC | 5.70 | 0.003 | 1.94 | 0.96 |
| 811.606 | PC | 5.97 | 0.002 | 2.34 | 1.23 |
| 814.62 | DGCC | -3.53 | 0.039 | 0.63 | -0.67 |
| 815.622 | | -3.59 | 0.037 | 0.63 | -0.67 |
| 832.562 | | 6.21 | 0.002 | 4.17 | 2.06 |
| 834.577 | HexCer | 3.69 | 0.033 | 1.75 | 0.80 |
| 838.634 | PC | 3.91 | 0.027 | 2.25 | 1.17 |
| 839.663 | | 7.41 | 0.001 | 2.88 | 1.53 |
| 840.666 | PC/DGTA/DGTS/DGCC | 7.71 | 0.001 | 3.33 | 1.74 |
| 845.701 | TG | -3.72 | 0.033 | 0.28 | -1.82 |
| 846.703 | HexCer | -3.76 | 0.033 | 0.22 | -2.16 |
| 848.557 | PC | 3.37 | 0.047 | 2.33 | 1.22 |
| 849.56 | Pyro b | 4.35 | 0.015 | 3.43 | 1.78 |
| 865.677 | MGDG | 6.33 | 0.002 | 2.41 | 1.27 |
| 867.694 | MGDG | 7.66 | 0.001 | 2.76 | 1.46 |
| 868.697 | DGTA/DGTS/DGCC | 7.87 | 0.001 | 2.97 | 1.57 |
| 869.708 | MGDG | 3.63 | 0.035 | 1.55 | 0.63 |
| 870.715 | DGTA/DGTS/DGCC | 3.56 | 0.038 | 1.63 | 0.71 |
| 873.69 | | -3.72 | 0.033 | 0.59 | -0.76 |
| 874.693 | | -3.33 | 0.048 | 0.56 | -0.84 |
| 876.584 | SHexCer | 3.68 | 0.033 | 2.78 | 1.48 |
| 881.676 | | 7.47 | 0.001 | 4.26 | 2.09 |
| 883.69 | PG/CerPE | 5.87 | 0.002 | 2.16 | 1.11 |
| 884.695 | DGCC | 7.23 | 0.001 | 2.43 | 1.28 |
| 897.74 | | 3.49 | 0.040 | 1.85 | 0.89 |
| 901.663 | | -4.71 | 0.010 | 0.49 | -1.04 |
| 902.667 | | -4.14 | 0.020 | 0.48 | -1.05 |

Table S14. T-test results of metabolites that were significantly different in relative intensities between SS8-corals under ambient versus elevated temperatures. A fold change (FC) > 1 indicates that this metabolite was enriched under ambient temperature.

| m/z | Annotation | T.stat | $P_{adj}$ | FC | Log2(FC) |
|---|---|---|---|---|---|
| 612.52 | | 4.57 | 0.043 | 4.05 | 2.02 |
| 626.534 | Cer | 4.59 | 0.043 | 3.65 | 1.87 |
| 654.567 | Cer | 5.02 | 0.031 | 3.23 | 1.69 |
| 748.57 | DGCC | 4.50 | 0.044 | 1.91 | 0.93 |
| 766.575 | PC/PE | 5.81 | 0.015 | 1.94 | 0.96 |
| 767.579 | PE | 5.62 | 0.015 | 1.92 | 0.94 |
| 772.572 | DGCC | 5.73 | 0.015 | 3.26 | 1.70 |
| 774.587 | DGCC | 7.57 | 0.004 | 3.94 | 1.98 |
| 775.591 | TG | 8.23 | 0.004 | 4.51 | 2.17 |
| 788.604 | PC | 4.90 | 0.032 | 2.89 | 1.53 |
| 810.599 | PC | 6.52 | 0.009 | 2.71 | 1.44 |
| 811.606 | PC | 4.40 | 0.047 | 2.06 | 1.04 |

Table S15. Details of all raw data and R codes for statistical analysis of this study (available on doi: 10.5281/zenodo.8008851).

| Category | File name | Description |
|---|---|---|
| Symbiodiniaceae community | 2021CoralHeatShock_ITS2_final for publication.html | R codes |
| | 2021CoralHeatShock_ITS2_metadata.csv | Metadata |
| | 2021CoralHeatShock_ITS2_profiles.absolute.abund_and_meta | Symportal output |
| | 2021CoralHeatShock_ITS2_seqs.absolute.abund_and_meta | Symportal output |
| | 2021CoralHeatShock_ITS2_seqs.fasta | Symportal output |
| Symbiodiniaceae community (year 1 and 2) | 2021CoralHeatShock_ITS2_Y1Y2_final for publication.html | R codes |
| | 2021CoralHeatShock_ITS2_Y1Y2_metadata.csv | Metadata |
| | 2021CoralHeatShock_ITS2_Y1Y2_profiles.absolute.abund_and_meta | Symportal output |
| | 2021CoralHeatShock_ITS2_Y1Y2_seqs.absolute.abund_and_meta | Symportal output |
| | 2021CoralHeatShock_ITS2_Y1Y2_seqs.fasta | Symportal output |
| Symbiodiniaceae cell density | 2021CoralHeatShock_CellCount_final-for-publication.html | R codes |
| | 2021CoralHeatShock_CellCountv2.csv | Raw data |
| Coral size and survival | 2021CoralHeatShock_Survival_final-for-publication.html | R codes |
| | 2021CoralHeatShock_Survival_final.csv | Raw data |
| Photochemical efficiency | 2021CoralHeatShock_iPAM-final-for-publication.html | R codes |
| | 2021CoralHeatShock_iPAMmerged.csv | Raw data |
| Metabolite profiles | 2021CoralHeatShock_MSI | Raw data |
| | All_files.slx | Raw spectra |
| | All_files.sbd | Raw spectra |

**SUPPLEMENTARY METHODS S1**

A pilot study was conducted to test the bleaching efficiency of various menthol concentrations (0.19, 0.38. 0.58 mM), combination with Diuron (50 µM) and coral fragment sizes (single polyp, ~4 polyps and 10 polyps). The treatment that removed the most native Symbiodiniaceae cells and resulted in the lowest coral mortality (i.e., 0.38 mM of menthol on ~4-polyp sized coral fragments) was selected for this study.

**SUPPLEMENTARY METHODS S2**

One mL of *C. proliferum* (WT10) and *Durusdinium* sp. culture at $10^4$ cell $mL^{-1}$ was sampled to make three 50:50 *Cladocopium*: *Durusdinium* mock communities to assess for potential sequencing bias. The *Durusdinium* culture (SCF 086.01) was originally isolated from *Porites lobata* from Davies reef, GBR. Only *Cladocopium* and *Durusdinium* were selected for the mock community as wild *G. fascicularis* only harboured these genera. DNA from the mock community cultures was extracted and amplified separately; their amplicons were quantified with PicoGreen and an equal amount from each culture was pooled to make the mock community.

**SUPPLEMENTARY METHODS S3**

**DNA extraction, PCR amplification and library preparation**

All samples collected for DNA metabarcoding were snap frozen in liquid $N_2$ and stored at -80°C until DNA extraction. Sample DNA was extracted using a salting-out method (Wilson et al. 2002) with the addition of a 15 min incubation with 10 mg $mL^{-1}$ lysozyme, and 20 s bead beating at 30 Hz with 100 mg of 425- 600 sterile acid-washed glass beads. Three extraction blanks were included. The Symbiodiniaceae ITS2 primers: Sym_Var_5.8S2 [5' GTGACCTATGAACTCAGGAGT**C**GAATTGCAGAACTCCGTGAACC 3'] (Hume et al. 2015); Sym_Var_Rev [3' CTGAGACTTGCACATCGCAGCCGGGTTCWCTTGTYTGACTTC ATGC 5'] (Hume et al. 2013) with Illumina adapters (underlined- specific for the Walter and Eliza Hall Institute (WEHI)) were used to amplify the partial 5.8S, entire ITS2 and partial 28S rDNA genes. PCR was performed in triplicate; each of the 15 µL reaction consisted of 7.5 µL of MyTaq HSRed MasterMix (Bioline, Australia), 1.5 µL of each primer (10 µM), 3.5 µL of nuclease-free water, and 1 µL of DNA template. Three no template controls were included. PCR was conducted in a thermal cycler (SimpliAmp$^{TM}$ Thermal Cycler, Thermo Fisher Scientific, Scoresby, Australia) under the following conditions: one initial denaturation cycle at 95 °C for 3 min, 18 amplification cycles (denaturation at 95 °C for 15 s, annealing at 55 °C for 30 s and extension at 72 °C for 30 s), one extension cycle at 72 °C for 7 min, and a final hold temperature of 4 °C.

Triplicate PCR products were pooled into 20 µL, and each product pool were purified using Ampure XP magnetic beads. The purified DNA was resuspended in 40 µL of nuclease-free water. Ten µL of DNA was combined with 10 µL of 2x Taq master mix (M0270S, New England

BioLabs,Notting Hill, Australia), 1 µL of forward indexing, and 1 µL of reverse indexing primers. PCR was conducted under: one initial denaturation cycle at 95 °C for 3 min, 24 amplification cycles (denaturation at 95°C for 15 s, annealing at 60°C for 30 s and extension at 72°C for 30 s), one extension cycle at 72°C for 7 min, and a final hold temperature of 4°C. Product quantity and size were visualized on 1% agarose 1 x TAE agarose gel. A total of 5 µL from each reaction was pooled and a final bead clean-up was performed on the 50 µL pooled volume. Illumina MiSeq v3 sequencing was conducted at WEHI.

### SUPPLEMENTARY METHODS S4

#### Mass spectrometry imaging

##### *Sample collection and fixation*

Coral polyps were sampled with a bone cutter. To minimize the presence of mucus that would hinder ionization, the polyps were left to rest in their aquaria for seven hours after cutting before fixation. Each polyp was allocated an individual well of a 12-well plate containing 2 mL of seawater and incubated in the dark at their respective temperatures (27°C or 32°C) for 20 min. The incubation period allowed the coral tentacles to be extended before anesthetizing by slowly adding 1 mL of 0.4 M $MgCl_2$. Samples were left in the dark for another 15 min, thereafter rinsed twice with Milli Q water, and carefully dried with a Kimwipe. The polyps were then embedded in a carboxymethyl cellulose (CMC)-gelatin mix (a ratio of 10 mL CMC + 0.123 g of gelatin) with a pair of forceps and snap-frozen on dry ice. To prepare the CMC-gelatin mix, 100 mL of CMC was warmed up in a glass bottle in a microwave at low heat (40% power, 10 s at a time x 3 times) until the bottle was warm to touch but not boiling. Then 1.353 g of gelatin was slowly added to the CMC in batches and mixed thoroughly with a spatula. Excessive stirring was avoided as this could create bubbles in the mixture. The CMC-gelatin mix was stored in ~25-27°C, covered with aluminum foil and used within 6 weeks.

##### *Cryosectioning and matrix spray*

Samples were cryosectioned in a cryostat (Leica CM 1860, -25°C) with a disposable high-profile diamond blade, following a method adapted from (Kawamoto and Kawamoto 2014, 2021; Wada et al. 2016; Boughton et al. 2020). The samples were mounted on the cryostat with optimal cutting temperature compound (OTC) and sectioned at 12 µm thickness until arriving to the middle of the animal, where maximum areas of the frozen body and tentacles were exposed. To increase structural integrity of the section, a thin layer (~2 µm) of CMC-gelatin mix was applied on the exposed surface of the sample each time before collection. The section was collected at 12 µm thickness on a cryo-film, thereafter freeze-dried under ~-50°C and ~0.08 mbar for 12 min. Two desiccated sections per sample were mounted on a stainless-steel sheet with carbon tapes and coated with a matrix (α-Cyano-4-hydroxycinnamic acid, HCCA) to assist ionization. A total of 30 mg of HCCA was dissolved in 6 mL of solvent (70% Acetonitrile, 30% H2O with 0.1% Trifluoroacetic acid) for each matrix spray run. Matrix coating was carried out with a HTX TM-

Sprayer™ at a flow rate of 70 µL $min^{-1}$ for six passes under a nozzle temperature of 75°C. The sections were desiccated for 20 min prior to imaging.

***MALDI-MSI analysis***

MALDI-MSI analysis was performed on a Bruker SolariX (7T XR hybrid ESI–MALDI–FT–ICR–MS) with a mass resolving power of 200000 and equipped with a SmartBeam II UV laser. The instrument was calibrated with a red phosphorus standard prior to data collection to ensure a mass error of < 1.5 ppm. Two technical replicates were imaged per sample and three samples were imaged per MSI run. FlexImaging 4.1 (Bruker Daltonics) was used to define the area of interest for imaging and data acquisition was controlled via Bruker Daltonics ftmsControl 2.1.0. Spectra were collected at a spatial resolution of 50 µm and a range of 150-2000 m/z under positive ion mode, with the laser diameter and power set to 45 µm 52%. A total of 500 laser shots were applied at each 50 µm pixel at a frequency of 2 kHz. Each section required ~4 h to image. SCiLS was used to combine the spectra files of all samples, remove noises (threshold cut off = 9000) and generate a m/z peak list. Any remaining background noises (e.g., instrument noises beyond the cut off, CMC/gelatin signals) were identified by creating a region of interest (ROI) in a known background area and removed from the peak list. Any remaining background noises (e.g., instrument noises beyond the cut off, CMC/gelatin signals) were identified by creating a region of interest (ROI) in a known background area and removed from the peak list. The final peak list consists of 418 metabolites, each was visually confirmed on SCiLS that it is associated with the biological tissues. The final peak list consists of 418 metabolites. The intensities of the metabolites (average peak area) were normalized with root mean square to account for potential intensity differences between MSI runs. As coral sections vary slightly in size, the total surface area of a section was used to normalize metabolite intensities.

### SUPPLEMENTARY RESULTS

While the ITS2 mock community samples consisted of 50:50 *Cladocopium*: *Durusdinium* amplicons, on average, their ITS2 profiles comprised of 36% *Cladocopium*, 52% *Durusdinium* and 12% non-profile sequences. This suggests a slight underestimation of the relative abundance of *Cladocopium* sp. presented in this study, meaning that the corals were likely to have had a slightly higher relative abundance in SS8 and WT10 than presented. Nevertheless, this slight underestimation does not change the interpretations of this study.

### References

Boughton BA, Thomas ORB, Demarais NJ, Trede D, Swearer SE, Grey AC (2020) Detection of small molecule concentration gradients in ocular tissues and humours. Journal of Mass Spectrometry 55:e4460

Hume B, D'Angelo C, Burt J, Baker AC, Riegl B, Wiedenmann J (2013) Corals from the Persian/Arabian Gulf as models for thermotolerant reef-builders: Prevalence of clade C3

Symbiodinium, host fluorescence and ex situ temperature tolerance. Marine Pollution Bulletin 72:313–322

Hume BCC, D'Angelo C, Smith EG, Stevens JR, Burt J, Wiedenmann J (2015) Symbiodinium thermophilum sp. nov., a thermotolerant symbiotic alga prevalent in corals of the world's hottest sea, the Persian/Arabian Gulf. Sci Rep 5:8562

Kawamoto T, Kawamoto K (2014) Preparation of thin frozen sections from nonfixed and undecalcified hard tissues using kawamot's film method (2012). In: Hilton M.J. (eds) Skeletal Development and Repair: Methods and Protocols. Humana Press, Totowa, NJ, pp 149–164

Kawamoto T, Kawamoto K (2021) Preparation of thin frozen sections from nonfixed and undecalcified hard tissues using Kawamoto's film method (2020). In: Hilton M.J. (eds) Skeletal Development and Repair: Methods and Protocols. Springer US, New York, NY, pp 259–281

Wada N, Kawamoto T, Sato Y, Mano N (2016) A novel application of a cryosectioning technique to undecalcified coral specimens. Mar Biol 163:117

Wilson K, Li Y, Whan V, Lehnert S, Byrne K, Moore S, Pongsomboon S, Tassanakajon A, Rosenberg G, Ballment E, Fayazi Z, Swan J, Kenway M, Benzie J (2002) Genetic mapping of the black tiger shrimp *Penaeus monodon* with amplified fragment length polymorphism. Aquaculture 204:297–309
